# A Data-Intelligence-Intensive Bioinformatics Copilot System for Large-scale Omics Researches and Scientific Insights

**DOI:** 10.1101/2024.05.19.594895

**Authors:** Yang Liu, Rongbo Shen, Lu Zhou, Qingyu Xiao, Jiao Yuan, Yixue Li

**Author notes:** These authors contributed to the work equally and should be regarded as co-first authors (Yang Liu, Rongbo Shen, Lu Zhou). Corresponding authors: Jiao Yuan, Yixue Li.

## Abstract

Advancements in high-throughput sequencing technologies and artificial intelligence offer unprecedented opportunities for groundbreaking discoveries in bioinformatics research. However, the challenges of exponential growth of omics data and the rapid development of artificial intelligence technologies require automated big biological data analysis capability and interdisciplinary knowledge-driven scientific insight. Here we propose a data-intelligence-intensive bioinformatics copilot (Bio-Copilot) system that synergizes AI capabilities with human expertise to facilitate hypothesis-free exploratory research and inspire novel scientific insights in large-scale omics studies. Bio-Copilot forms high-quality intensive intelligence through close collaboration between multiple agents, driven by large language models (LLMs), and human experts. To augment the capabilities of Bio-Copilot, this study devises an agent group management strategy, an effective human-agent interaction mechanism, a shared interdisciplinary knowledge database, and continuous learning strategies for the agents. We comprehensively compare Bio-Copilot against GPT-4o and several leading AI agents across diverse bioinformatics tasks, using a broad range of evaluation metrics. Bio-Copilot achieves the overall state-of-the-art performance across all tasks, while showcases exceptional task completeness. Furthermore, in the application of constructing a large-scale human lung cell atlas, Bio-Copilot not only reproduces the intricate data integration process detailed in a seminal study but also introduces a hierarchical annotation strategy to capture the continuous nature of cellular states and uncovers the characteristics of rare cell types, highlighting its potential to unravel hidden complexities in biological systems. Beyond the technical achievements, this study also underscores the profound implications of integrating AI capabilities with expert knowledge in accelerating impactful biological discoveries and exploring uncharted territories in life sciences.

## I. Introduction

### A. Background and Significance

Bioinformatics, as an interdisciplinary field that marries biology, computer science, mathematics, and statistics, has experienced unprecedented growth over the past few decades^1,2^, driven primarily by advancements in high-throughput sequencing technologies^3,4^ such as genomics^5,6,7,8^, transcriptomics^9,10^, proteomics^11,12,13^ and metabolomics^14,15^. With the plummeting costs and the accelerated proliferation of these technologies, the amount of biological data being generated has exploded exponentially, presenting a wealth of opportunities for groundbreaking discoveries^16,17^. This torrential influx of data has fundamentally revolutionized our understanding of life sciences, ranging from fundamental molecular biology^18^ to practical clinical applications and the burgeoning domain of personalized medicine^19,20^.

In recent years, the scientific research landscape in bioinformatics has been undergoing a profound shift^21,22^, propelled by the confluence of two major factors: the exponential growth of complex and interconnected biological data and the rapid advancement of artificial intelligence (AI) technologies. Despite the significant strides achieved in merging bioinformatics with AI technologies, current bioinformatics research still faces several critical challenges^23,24^. Firstly, integrating the strengths of AI with big biological data to propel advancements in bioinformatics research constitutes an ongoing and arduous pursuit^25,26^. Many researchers in the field are limited by a single area of expertise, but lack interdisciplinary knowledge and the ability to conduct cross-disciplinary scientific research. The effective application of AI in bioinformatics often requires a deep understanding of both the biological domain knowledge and advanced computational techniques, therefore there is a need for enhanced interdisciplinary collaboration to bridge the domain gap and unlock the AI-driven analytical capabilities for life science research^27^. Secondly, the rapid accumulation of high-dimensional and multimodal biological data, characterized by large-scale, complexity, diversity, noise, and contextual dependencies, necessitates the development and application of advanced AI technologies. AI must not only enable scalable, adaptable, and interpretable processing of big biological data, but also promote synergistic integration with expert knowledge, thereby enhancing overall intelligent capacity.

### B. Research Statement

In light of the revolutionary progress in AI technology and the exponential growth of biological data, this study delineates a data-intelligence-intensive research paradigm, as illustrated in Figure 1a. This paradigm integrates AI capabilities with human expertise in a cohesive collaboration to form intensive intelligence^28^, that leverages big biological data to propel hypothesis-free exploratory research driven by a data-intelligence-intensive methodology. This novel research paradigm aligns with the future trend of bioinformatics research, characterized by data-intensive and computation-intensive^29^ in big biological data processing, and knowledge-intensive and intelligence-intensive^30,31^ in the convergence of interdisciplinary. Within this paradigm, human expertise continually enhances and refines AI agents, while AI agents driven by large language models (LLMs), in turn, leverages multidisciplinary knowledge fusion and big biological data mining to generate novel insights, thereby improving our understanding of life sciences. By leveraging their complementary strengths, bioinformatics research gains the ability to traverse uncharted territories of knowledge, generating testable hypotheses, delineating novel research directions, and refining experimental designs to accelerate discoveries, thereby revealing previously unexplored associations and emergent characteristics within complex biological systems.

**Figure 1.**
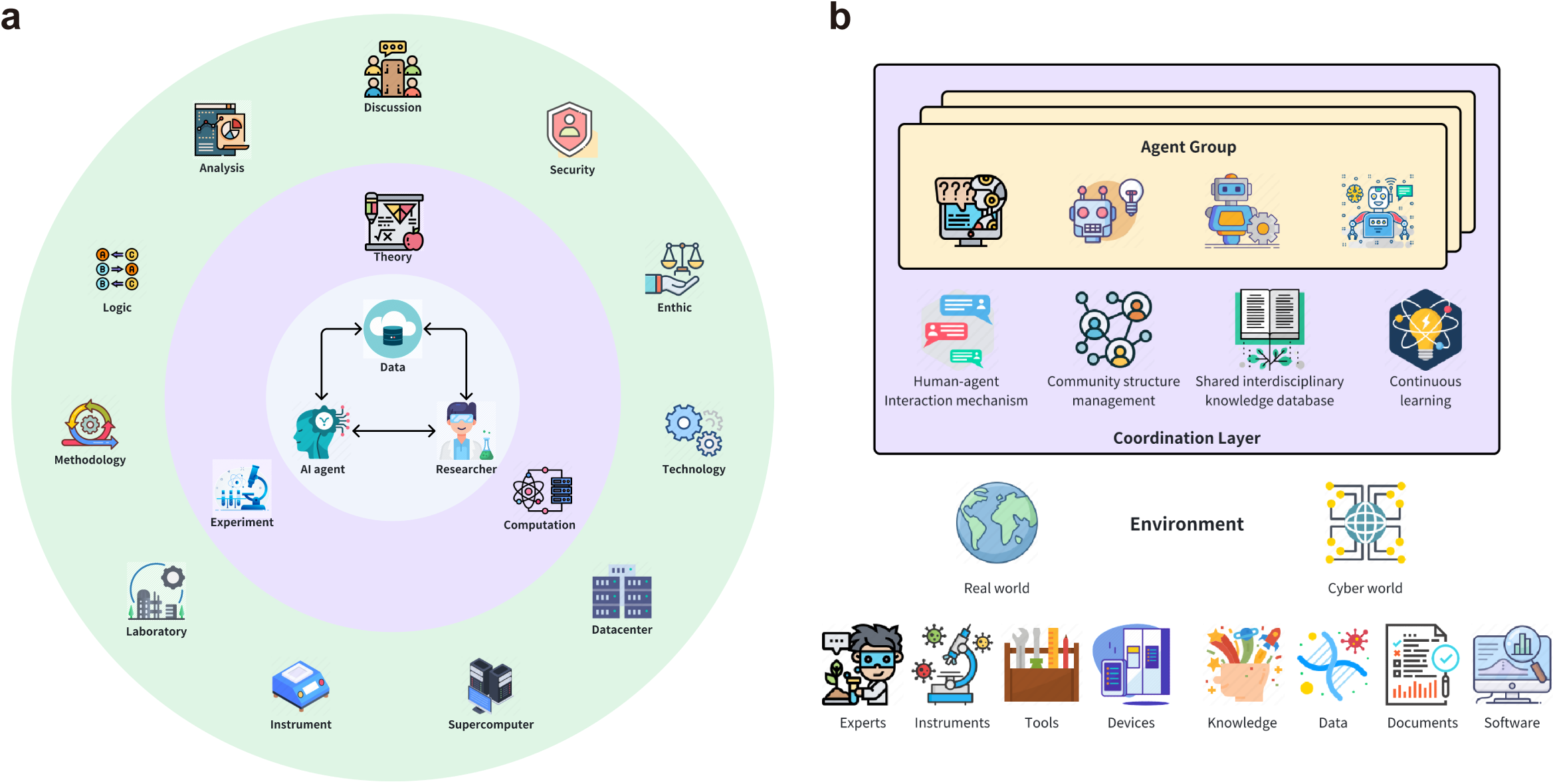
**a. Data-intelligence-intensive research paradigm**. Synergizing human expertise with AI capabilities forms intensive intelligence, that leverage big biological data to propel hypothesis-free exploratory research driven by a data-intelligence-intensive methodology. **b. The architecture of Bio-Copilot**. It comprises multiple agents with di-verse roles, a coordination layer, and an environment.

The data-intelligence-intensive research paradigm presents a promising perspective for addressing contemporary challenges in current bioinformatics. It embeds knowledge from diverse scientific disciplines into AI agents, who, as integral members of intensive intelligence, actively participate in bioinformatics researches, facilitating interdisciplinary collaboration and overcoming long-standing knowledge barriers. Moreover, this novel research paradigm, leveraging the complementary integration of expert knowledge with AI capabilities^32,33^, accelerates bioinformatics workflows by feature extraction from big biological data, proposing innovative scientific hypotheses, optimizing research designs, orchestrating computational resources for high-performance computing, and instituting fair and objective evaluation metrics.

In the data-intelligence-intensive research paradigm, both human experts and AI agents are core participants in scientific investigations. Here, we propose a data-intelligence-intensive bioinformatics copilot (Bio-Copilot) system that synergizes AI capabilities with human expertise to facilitate hypothesis-free exploratory research and inspire novel scientific insights in large-scale omics. Bio-Copilot utilizes the potent natural language processing capabilities of LLMs^34,35,36^ and their broad perspective on scientific issues to enable interdisciplinary knowledge embedding, interaction with human experts, inter-agent communication, and agent self-reflective learning, thereby forming high-quality intensive intelligence through close collaboration between multiple agents and human experts. Initiated by scientific insights from LLMs, Bio-Copilot first decomposes a bioinformatics task into modular, hierarchical steps, then configures agent groups and specifies the roles of agents within each group according to step characteristics, alongside formulating rules for task allocation and scheduling. Thereafter, researchers collaborate with the planning agent groups to generate more explicit execution plans for the respective steps. Finally, according to the generated execution plans, the action execution agent groups proceed gather necessary resources from the environment and sequentially accomplish each execution plan.

In the context of data-intelligence-intensive research paradigm, this study uses Bio-Copilot to substantiate the following scientific hypothesis: Firstly, the workflows planned by high-quality intensive intelligence, involving collaboration between researchers and multiple agents in Bio-Copilot, can enable bioinformatics researches to achieve superior results, significantly improving research efficiency and overcoming the limitations^37^ inherent to standalone LLM applications. Secondly, the interactions between researchers and Bio-Copilot can inspire novel scientific insights, mitigate comprehension biases in LLMs pertaining to prompts, context, and both long-term and short-term memory, and address the problems of incomplete information during action execution, thereby enabling adaptability to complex bioinformatics research. Thirdly, strengthened by multidisciplinary knowledge empowerment and continuous learning of its agents, Bio-Copilot can make better decisions, plan better workflows and select optimal tools for bioinformatics research.

### C. Main Contributions

The main contributions of this study are enumerated as follows:

- This study proposes a data-intelligence-intensive Bio-Copilot system, which forms high-quality intensive intelligence through close collaboration between multiple agents and human experts to facilitate hypothesis-free exploratory research and inspire novel scientific insights in large-scale omics.
- To augment the capabilities of Bio-Copilot, this study devises an agent group management strategy, an effective human-agent interaction mechanism, a shared interdisciplinary knowledge database, and continuous learning strategies for agents.
- This study systematically compares Bio-Copilot against GPT-4o and several AI agents in diverse bioinformatics tasks using a broad range of evaluation metrics, and demonstrates the advancement of Bio-Copilot.
- Through the application of Bio-Copilot in constructing a large-scale human lung cell atlas, this study demonstrates that Bio-Copilot can not only reproduce existing workflow, but also carry out exploratory research tasks.

## II. Literature Review

### A. The development trend of bioinformatics

The evolution of bioinformatics, tracing back to the mid-20th century, has lately entered into a phase of accelerated development, catalyzed by the successful completion of the Human Genome Project^38,39,40,41^ and the groundbreaking advancements in high-throughput sequencing technologies^3,4,42,43^. The explosive growth of high-throughput sequencing data, exemplified by genomics^5,6,7,8^, transcriptomics^9,10,44^, proteomics^11,12,13^, and metabolomics^14,15^, has provided unprecedented data resources for life sciences research. Driven by the data-intensive research paradigm, researchers developed more efficient methods for processing and analyzing large-scale biological datasets to elucidate the structures and functions of biomolecules, cells, tissues, organs, and entire organisms, encompassing gene regulatory networks^45,46^, epigenetic modifications^47,48^, cellular heterogeneity^49,50^, evolutionary relationships^51,52^, and disease mechanisms^53,54,55^. In precision medicine^19,20,56^, the integration of genomic, cellular phenotype, and clinical data enabled multi-modal data analyses^57,58^, advancing applications in disease prevention, diagnosis, personalized treatment planning, and drug response prediction. Recently, with the breakthroughs in AI technology, machine learning and deep learning methods have become paramount tools in large-scale biological data analysis, excelling in handling complex nonlinear relationships and demonstrating substantial potential in microscopy image analysis^59,60^, protein structure prediction^61,62,63,64^, deciphering gene regulatory networks^45,46,65,66^, biomarker detection for diseases^67^, diagnostic and prognostic assessments^68^, and drug discovery and design^69,70,71^.

To address the mounting volume of biological data and comprehensively unravel complex life systems, the progression of bioinformatics evolved advancements in technology, computational methodology innovation, and interdisciplinary collaboration, propelling the development of life sciences through deeper analysis and understanding of biological data. For instance, with the maturation and cost reduction of single-cell sequencing technologies, single-cell multi-omics^72,73^ studies integrated multi-omics data in spatiotemporal dimensions, elucidating the fundamental laws governing life systems across different molecular levels (i.e., genetic, transcriptional, proteomic, and metabolic), thereby enabling a finer dissection of cellular heterogeneity, intercellular interactions, and dynamic cellular processes, which fostered novel discoveries in precision medicine, tumor biology, and developmental biology. Third-generation sequencing technologies^74^, with their high sensitivity, long reads, and real-time sequencing capabilities, generated full-length transcript sequencing data, facilitating rapid epigenetic modification identification, and contributing to the study of alternative splicing^75^ and non-coding RNA structures^76,77,78^, thereby enhancing our understanding of gene transcription regulation, non-coding RNA biology, and the mechanisms underlying disease development. AI technologies are becoming increasingly embedded in bioinformatics research, capable of handling complex pattern recognition, big data analytics, and predictive modeling, enhancing accuracy and efficiency from omics data analysis to protein structure prediction, to disease diagnostics and drug development. To enhance the efficiency and reproducibility of bioinformatics research, automated workflows provided standardized data processing, elastic resource allocation with distributed computing, and workflow management and monitoring, thereby reinforcing the rigor and objectivity of scientific investigations.

### B. Large Language Models

Large language models (LLMs) are deep learning architectures based on the Transformer^79^, featuring stacked multi-layered structures and positional encoding, which leverage self-attention mechanisms to process input text sequences in parallel, thereby efficiently capturing long-range dependencies and learning linguistic structures and patterns. LLMs typically undergo pre-training on vast amounts of unlabeled text data, acquiring general linguistic patterns and structures. Following the pre-training phase, LLMs can be fine-tuned to specific tasks such as question answering, text classification, or text generation, tailored to the precise requirements of the given application. LLMs, with their massive parameters and deep network architectures, excel at understanding complex contextual relationships and generating coherent, high-quality text^80^. With technological progress, some LLMs incorporate multimodal learning capabilities, enabling comprehension of data such as images and audio in addition to text, and thereby achieving cross-modal information processing and synthesis.

The proposal of Transformer^79^ architecture in 2017 marked the dawn of the era of LLMs. In 2018, the unveiling of BERT^81^ by Google and GPT-1^82^ by OpenAI introduced pre-training techniques, which substantially improved the performance capacities of LLMs. The successive releases of the GPT series^83,84^ propelled the rapid advancement of LLMs, with model parameters swiftly surging into trillions. Particularly, ChatGPT^83^ and GPT-4^84^ demonstrated unprecedented capabilities in language generation and comprehension, achieving remarkable performance across diverse applications such as question answering, text composition, code generation, and conversational system^85,86,87^. Subsequently, LLMs integrated multimodal processing capabilities, enabling joint modeling of diverse data types such as images and text, and thereby offering enhanced visual generation abilities (e.g., Sora^88^).

While LLMs have demonstrated remarkable capabilities, they also come with inherent limitations^37^. The output of LLMs is highly contingent upon training data, thereby inheriting biases, factual inaccuracies, and illusion issues from biased, erroneous, or incomplete training datasets. While capable of handling and generating sophisticated language structures, LLMs, as statistical prediction engines, lack the comprehension, common sense, autonomous planning, and self-learning capabilities inherent to humans, rendering them potentially inadequate in tasks requiring profound understanding or logical reasoning. Owing to the complexity of LLMs, the decision-making processes of LLMs tend to be opaque, rendering explanations for specific predictions or generations elusive.

### C. AI Agent

The origins of AI agent development can be traced back to the emergence of expert systems, which leveraged rule-based knowledge databases to mimic human experts in solving domain-specific problems. Subsequently, reinforcement learning^89,90^ emerged as a vital machine learning paradigm integrated into AI agents, offering a pathway for AI agents to learn through iterative interactions with their environments. In recent years, the revolutionary advancements in deep learning technologies, highlighted by the introduction of LLMs such as the GPT series, empowered LLM-based agents with formidable natural language processing capabilities. Agents^91,92,93,94^, capable of autonomously perceiving their environment, cognitive reasoning, decision-making, and executing actions through tool invocation, emerged as a highly promising direction in the pursuit of general artificial intelligence. Specifically, an AI agent comprises four modules: the perception module for gathering environmental information, the cognition and decision-making module to analyze inputs and devise action strategies, the memory module to archive knowledge and past behaviors, and the action module to implement decisions by manipulating tools to impact the environment. AI Agents execute tasks through a perceptual-decisional-action loop, subsequently engaging in reflective evaluation of the results, which informs learning to acquire knowledge, stored in the memory module, thereby optimizing future decision-making and actions. Agents demonstrate human-like intelligence behavior^95^ through autonomous environmental perception, decision-making process, and an aptitude for learning and adapting via dynamic environmental engagements.

Prominent existing single-agent frameworks such as AutoGPT^96^, BabyAGI, and HuggingGPT^97^ facilitated task planning and tool invocation, thereby accomplishing user-defined objectives effectively. For instance, the Coscientist^98^ system, powered by GPT-4, autonomously conducted chemical research, involving automated experiment planning and execution, along with predicting and refining reaction pathways through sophisticated modeling capabilities. Researchers interacted with the Coscientist system to accomplish six pivotal tasks: planning synthesis routes of known compounds using public data, efficiently retrieving and navigating hardware documentation, executing advanced directives in cloud labs, precisely controlling liquid handling equipment, multitasking across multiple hardware modules for complex operations, integrating diverse datasets and analyzing past experimental data for optimization problem-solving. For another example, the ChemCrow^99^ system, driven by LLMs, autonomously planned and executed tasks including the synthesis of organic compounds, drug discovery, and material design. By incorporating tools such as web searches, literature exploration, and interactive coding interfaces, the ChemCrow system augmented its capability to acquire and comprehend external information, thereby showcasing the potential for enhanced scientific task management. In the field of computational pathology, PathChat^100^, a multimodal generative AI Copilot system, demonstrated the ability to combine histological images, clinical data, and subsequent immunohistochemical (IHC) results for interactive differential diagnosis in unknown primary cancer cases, aiming to assist pathologists improving the accuracy and efficiency in diagnosis and understanding complex cases. PathChat was designed to handle pathology-related queries in multiple-choice diagnostic questions, demonstrating its potential in pathology diagnostic support. While LLMs have propelled the evolution of AI agent technologies, single-agent systems still grapple with inherent limitations. Firstly, the capabilities of AI agents are confined by the prowess of their underlying LLMs, which can lead to biased, hallucinatory, erroneous, or outdated action decisions during complex tasks, often due to the quality of training data. When dealing with extensive texts, iterative conversations, or complex contexts, AI agents may misconstrue user intentions and offer partially relevant responses, hampered by the finite memory retention of LLMs and limitations in comprehending prompt cues and contextual associations. Secondly, while individual agents can adopt multiple roles, they often rapidly converge onto a specific viewpoint influenced by prompts or earlier conversation turns, thereby missing out multiperspective and holistic abilities for problem analysis. Lastly, single-agent systems typically decompose tasks into linear workflows, resulting in inefficient execution and inadequate responsiveness to dynamically changing environments.

LLM-based single-agent systems focus on internal reasoning and decision-making processes as well as interactions with the external environment. Inspired by human collaboration in tackling complex tasks, LLM-based multi-agent systems emphasize diverse agent groups, inter-agent interactions, and collective decision-making protocols. Compared to single-agent systems, multi-agent systems orchestrate a consortium of diverse agents with distinct capabilities and roles, simulating intricate real-world scenarios via inter-agent cooperation, harnessing the collective intelligence born of their individual strengths to tackle more dynamic and intricate tasks such as software development, scientific experiment, and embodied AI.

Recently, the open-source community witnessed the rise of several prominent multi-agent frameworks^101,102,103^, such as MetaGPT^104^, CAMEL^105^, and AutoGen^106^. MetaGPT embeded human workflow procedures into the actions of LLM-based agents, employing structured outputs and a publish-subscribe mechanism, effectively resolving information redundancy in multi-agent conversations and mitigating the prevalence of illusions in complex tasks. CAMEL leveraged prompt engineering techniques to delineate the roles of agents, facilitating autonomous collaboration between them, and directing agent conversations to progress in alignment with mission objectives. AutoGen emerged as a highly customizable multi-agent framework, enabling researchers to customize the LLMs of agents, communication protocols and workflow management, thereby tackling intricate tasks. However, multi-agent systems also face some challenges. Fundamentally, the design and implementation of multi-agent systems entail heightened complexity, requiring meticulous attention to issues of inter-agent communication, collaboration, and conflict resolution mechanisms. Additionally, multi-agent systems encounter sophisticated interaction patterns in the dynamic contexts, which can lead to information overlaps and non-productive discourse, ultimately hindering the effectiveness of task execution. Furthermore, multi-agent systems are characterized by significant computational resource consumption and a heightened cost associated with customization.

## III. Results

### A. Architecture and advancements of Bio-Copilot

#### Architecture

The architecture of Bio-Copilot comprises multiple agents with diverse roles, a coordination layer, and an environment, as illustrated in Figure 1b. Agents, the core components of Bio-Copilot, employ LLMs as their controlling nucleus and exhibit a high degree of autonomous capability. They are capable of perceiving environmental information, deciphering complex natural language instructions (such as the prompts from experts in the environment), conducting cognitive reasoning, and executing actions through tool invocations. Agents can act various roles with specific responsibilities, such as information acquisition, task planning, code generation, task execution, and etc. The coordination layer serves as a bridge for agents to perceive environmental information and feedback to influence the environmental state, while harmonizing the behaviors of agents such as task planning and allocation, resource scheduling, conflict resolution, and collaborative strategies. Within the coordination layer, we devise an agent group management strategy, an effective human-agent interaction mechanism, a shared interdisciplinary knowledge database, and continuous learning strategies for agents, thereby augmenting the capabilities of Bio-Copilot. The environment, as an external context of Bio-Copilot, influences the behavior of agents, while the actions of these agents in turn modify the state of the environment. Specifically, the environment comprises elements from both the real world, such as experts, instruments, devices and tools, and the virtual world, including datasets, documents, knowledge databases, and software.

#### Advancements

Bio-Copilot employs an innovative design approach that emphasizes interdisciplinary collaboration, shared knowledge resources, and continuous interactive feedback, devising novel strategies in efficient human-agent interaction, agent group management, interdisciplinary knowledge empowerment, and continuous learning. Firstly, to address the complexity of bioinformatics problems, we devise user-friendly, highly real-time human-agent interaction mechanism, aimed to alleviate the misunderstandings of agents related to prompts, context, and both long-term and short-term memory, alongside issues of incomplete information during action execution. This human-agent interaction mechanism also supports natural language interactions and immediate feedback between human experts and Bio-Copilot, thereby inspiring new scientific insights. As a result, human experts can direct the activities of Bio-Copilot in real time and promptly assimilate analytical insights from Bio-Copilot. Secondly, to enhance collaboration efficiency, we devise agent group management strategies which can reduce redundant information exchange and inconsequential interactions to ensure efficient and accurate information flow. Thirdly, acknowledging the interdisciplinary nature of bioinformatics, we establish a shared knowledge database which encodes domain-specific expertise from diverse fields into a format intelligible within Bio-Copilot. Empowered by interdisciplinary knowledge, agents can dynamically retrieve relevant information from the shared knowledge database to formulate rational plans for task-specific requirements, thereby mitigating biases and illusions prevalent in LLMs during reasoning and decision-making. Lastly, to swiftly accommodate emerging and challenging bioinformatics research tasks, we devise continuous learning strategies for agents. Specifically, the multiple agents engage in both self-reflective learning^107^ from their own behaviors and competitive knowledge acquisition with fellow agents, progressively refining their decision-making and execution capabilities over time to enhance intensive intelligence. The aforementioned innovative design enhances Bio-Copilot in terms of language intelligence, social intelligence, knowledge empowerment, and learning intelligence.

### B. Benchmarking Bio-Copilot for the construction of a large-scale human lung cell atlas

#### Workflow description

In recent years, a series of single-cell sequencing studies dissected molecular profiles of cells in both normal and diseased tissues, and the rapidly advancing single-cell sequencing technology made it feasible to establish human cellular atlas (HCA) ^108^, which was listed among the seven notable technologies in 2024 by the scientific journal Nature^109^. A comprehensive cellular atlas of human lung tissue systematically unveiled molecular distinctions among various pulmonary cell populations, poised to revolutionize our understanding of cells in healthy and diseased lungs, thereby offering crucial insights for elucidating respiratory disease pathogenesis, vaccine development, immunotherapy, and tissue regeneration. Several independent studies embarked on constructing a cellular atlas for lung tissue. However, limitations in experimental techniques, sequencing methodologies, sample selection and cell quantity constrained comprehensive and unbiased capture of the diversity and heterogeneity inherent to lung tissues. Consequently, integrating datasets from these independent studies in an unbiased manner to construct a comprehensive cellular atlas of lung tissue would address these limitations and further facilitate the identification of variability patterns across different demographics, such as age, gender, BMI, smoking status, and etc.

In this study, we employ Bio-Copilot to construct a large-scale human lung cell atlas, thereby evaluating the capability of Bio-Copilot in handling data-intelligence-intensive bioinformatics research tasks. Referencing the HLCA^110^ study, the workflow is collaboratively generated by researchers and Bio-Copilot, which mainly comprises four tasks: dataset integration task, hierarchical clustering task, cell type annotation task, and downstream analysis task. Bio-Copilot adopts a consistent procedure for each task, comprising coarse-level planning, fine-level planning, and action execution, the example procedure for hierarchical clustering task is illustrated in Figure 2. Specifically, for each task objective, Bio-Copilot employs coarse-level planning group to formulate an initial plan, which then iteratively refined by fine-level planning group for each step. During the action execution of plan, researchers can intervene via human-agent interaction to set parameters and correct any erroneous actions and behaviors, as illustrated in Figure 2. Comprehensive workflow and implementation details for constructing the large-scale human lung cell atlas are in the Experimental setup section of Methodology.

**Figure 2.**
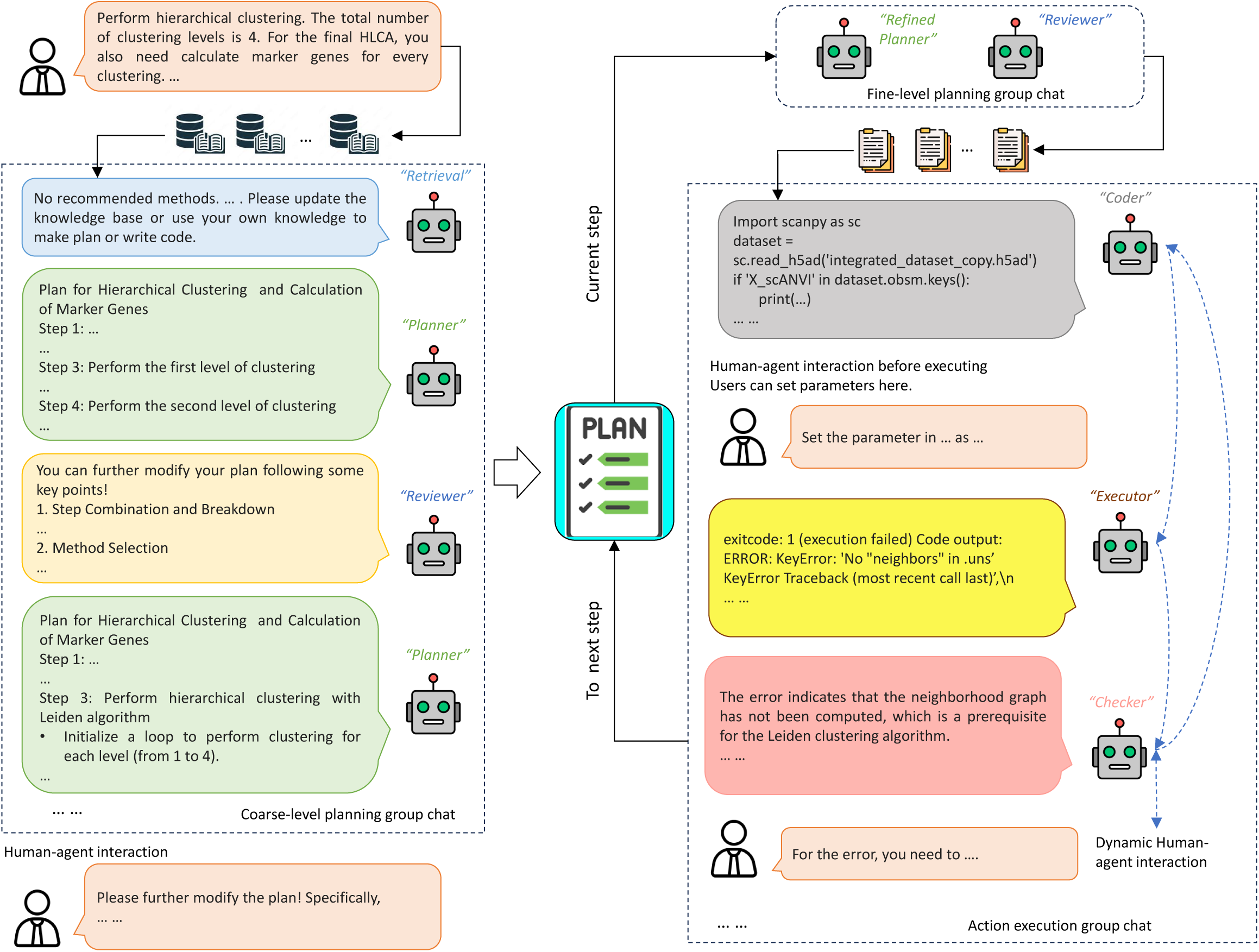
The workflow of Bio-Copilot. Bio-Copilot adopts a consistent procedure for each task, comprising coarse-level planning, fine-level planning, and action execution. The procedure involves close collaboration between multiple agents with human experts.

#### Performance evaluation

To fully evaluate the performance of Bio-Copilot, we systematically compare Bio-Copilot against the control group and the ablation group in all bioinformatics tasks in the workflow of constructing the large-scale human lung cell atlas, using a broad range of evaluation metrics. The control group comprises GPT-4o, AutoBA and AutoGen, where GPT-4o is a state-of-the-art LLM, AutoBA is an influential single AI agent for bioinformatics, and AutoGen is an influential multiple agent system. The ablation group comprises the exclusion of knowledge empowerment, self-reflective learning and document retrieval for Bio-Copilot, respectively. The evaluation of Bio-Copilot focuses on the following three aspects: task planning, code quality, and execution efficiency. We use an overall evaluation formula to synthesize all the evaluation aspects. Obviously, Bio-Copilot achieves the overall state-of-the-art performance in all tasks (Figure 3a), while showcases highest task completeness, as illustrated in Supplementary Figure 3a.

**Figure 3.**
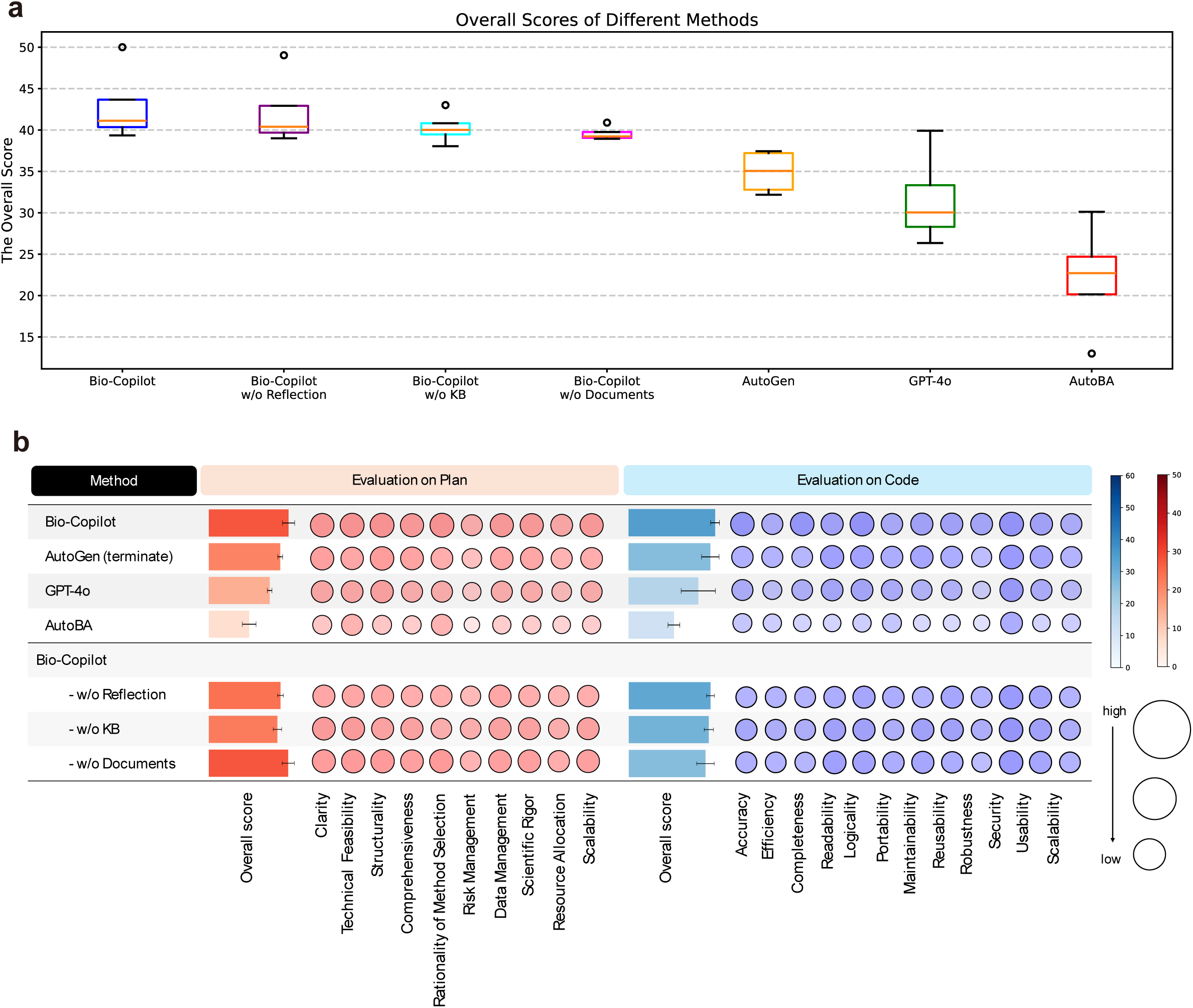
**a. Overall benchmarking results.** A overall evaluation formula is used to synthesize the task planning evaluation, code quality evaluation and execution efficiency evaluation for all methods. **b. The results of task planning evaluation and code quality evaluation.** In the task planning evaluation, GPT-4 employs 10 evaluation metrics to score the execution plans generated by all methods. In the code quality evaluation, GPT-4 employs 12 evaluation metrics to score the codes generated by all methods.

The task planning evaluation assesses the capabilities of Bio-Copilot in decomposing bioinformatics tasks and formulating execution plans. The evaluation metrics includes clarity, technical feasibility, structurality, comprehensiveness, rationality of method selection, risk management, data management, scientific rigor, resource allocation, and scalability. To ensure objectivity in this comparison, we employ GPT-4 to score the execution plans generated by each method, where the prompt of GPT-4 is illustrated in Supplementary Figure 1a. Obviously, Bio-Copilot achieves the overall state-of-the-art task planning performance in all tasks, as illustrated in Figure 3b. The execution plans generated by Bio-Copilot exhibit higher implementability in bioinformatics research tasks, compared to GPT-4o, AutoBA and AutoGen in the control group. It is further substantiated that intensive intelligence through close collaboration between multiple agents and experts facilitates the design of excellent scientific research programs. In addition, the comparison with the ablation group also demonstrates that knowledge empowerment, self-reflective learning and document retrieval can improve the task planning performance of Bio-Copilot. More scores of evaluation metrics for task planning are illustrated in Supplementary Figure 2a-b.

The code quality evaluation assesses the ability of Bio-Copilot to understand the execution plans. The evaluation metrics includes accuracy, efficiency, completeness, readability, logicality, portability, maintainability, usability, reusability, robustness, security, and scalability. This comparison also employs GPT-4 to score the quality of code generated by each method, where the prompt of GPT-4 is illustrated in Supplementary Figure 1b. Similarly, Bio-Copilot achieves the overall state-of-the-art code quality performance in all tasks, evidencing a more comprehensive understanding of execution plans, as illustrated in Figure 3b. The comparison with the control group demonstrates that Bio-Copilot can generate higher quality code for the execution plans. The comparison with the ablation group also demonstrates that knowledge empowerment, self-reflective learning and document retrieval are beneficial for Bio-Copilot in code generation. More scores of evaluation metrics for code quality are illustrated in Supplementary Figure 2c-d.

The execution efficiency evaluation assesses Bio-Copilot in task completeness, user interventions, runtime, computing resource cost, as illustrated in Supplementary Figure 3a-d. Obviously, Bio-Copilot can complete all tasks according to the generated codes, whereas methods in the control group fail to complete most of the tasks. More features of methods in the control group are illustrated in Supplementary Figure 3e. The user interventions represent the frequency with which the user directs the model. Bio-Copilot accomplishes all tasks with few user interventions, highlighting a high level of automation. Notably, AutoBA has no user intervention during execution, but it has a low task completeness, suggesting that proper user interventions during the execution of complex tasks can prevent task interruptions or failures due to misunderstanding or inadequate information. GPT-4o and AutoGen have about the same number of user interventions as Bio-Copilot across all tasks, but their tasks are not completed. This also substantiates the efficacy of our designed human-agent interaction mechanism, enabling Bio-Copilot to tackle complex bioinformatics tasks. The comparison with the ablation group demonstrates that knowledge empowerment, self-reflective learning and document retrieval can effectively reduce the user interventions. Across all tasks, Bio-Copilot has less overall rumtine than GPT-4o and AutoGen, while achieving a higher task completeness. Although AutoBA has less overall runtime, it has the lowest task completeness. In terms of resource consumption, the computing resource consumption of each method is within an affordable range, where 128 CPU cores, 32GB memory, and a GPU with 24GB memory can meet the usage requirements of Bio-Copilot.

Details on the evaluation metrics and the GPT-4 assessment approach (Supplementary Figure 1a-b) are provided in the Evaluation Metrics section of Methodology. The prompts of Bio-Copilot with self-reflective learning, knowledge empowerment and document retrieval are illustrated in Supplementary Figure 4a, 5a and 6a, respectively. In contrast, the prompts of the exclusion of self-reflective learning, knowledge empowerment and document retrieval in the ablation group are illustrated in Supplementary Figure 4b, 5b and 6b, respectively.

### C. Result analysis of the human lung cell atlas

#### Data integration and hierarchical clustering

Conforming to the workflow of constructing the large-scale human cell atlas, all tasks are performed by Bio-Copilot with expert collaboration. Bio-Copilot first performs the data integration task on multiple independent datasets. Utilizing the scANVI^111^ algorithm, Bio-Copilot accomplishes dimensionality reduction to embed cells from these datasets into a unified feature space. The scANVI algorithm mitigates batch effects arising from variations in sampling techniques and sequencing technologies across datasets, enabling cells of the same type to cluster in the feature space, thereby facilitating downstream analyses and visualizations.

Subsequently, Bio-Copilot conducts the hierarchical clustering task on the embeddings of the integrated dataset to identify distinct cell populations. By clustering cells in a hierarchical manner from coarse to fine resolutions, this approach enables researchers to identify cell populations at various levels and decipher the intricate relationships between them. Specifically, Bio-Copilot performs a four-level hierarchical clustering to categorize all cells within the dataset into distinct clusters at each level. The UMAP visualizations of the hierarchical clustering results from level 1 to level 4 are illustrated in Supplementary Figure 7a. These visualizations reveal that Bio-Copilot successfully reproduces the data integration task and hierarchical clustering task as reported in the original study. Furthermore, the batch effects between originally independent datasets are effectively mitigated after integration, in which most cells are grouped into a few of cell populations, and very few cells are isolated.

More details of Bio-Copilot performing the data integration task and hierarchical clustering task refer to the Experimental Setup section in Methodology.

#### Cell type annotation

In the original hierarchical annotations of these datasets, a considerable number of cells show unfilled categories (prefixed with numbers and inherited from upper levels during preprocessing) from level 2 to level 4, evidencing incomplete cell annotations under the original cell type nomenclature. Therefore, to refine the cell atlas and harmonize the cell type nomenclature, Bio-Copilot performs the cell type annotation task to re-annotate all clusters from level 2 to level 4. Specifically, Bio-Copilot ingests a database of human cell marker genes, and subsequently invokes the Decoupler^112^ tool to identify cell types for each cluster based on that database. This method of invoking tools is more stable and reproducible than directly leveraging LLMs for annotations^113^. The re-annotated cell types from level 2 to level 4 exhibit a stable hierarchical structure and favorable lineage characteristics, as illustrated in Figure 4a. This also underscores the continuity in cellular states within cell atlases, rendering a hierarchical annotation strategy for cell atlas construction facilitates lineage tracing and the discovery of novel subpopulations^114^, whereas single-cluster definitions inadequate for fully capturing the complexity of cell types. Due to the limited capabilities of the Decoupler tool and the database of human cell marker genes, a few cell clusters (a total of 4,579 cells, about 0.8% of the integrated dataset) cannot be identified as reliable cell types, so they are classified as unknown cell types. Subsequently, Bio-Copilot calculated the distribution of cells within each well-annotated cell type at level 4 across the original independent datasets, alongside entropy values, as illustrated in Figure 4b. From these computations, it is evident that the majority of cell types comprise cells from multiple datasets, whereas a few cell types originate predominantly from individual datasets. This again demonstrates that the batch effects between originally independent datasets are effectively mitigated in previous data integration task. Bio-Copilot further performs differential expression analysis and correlation analysis for the re-annotated integrated dataset at level 4, as illustrated in Supplementary Figure 8a. These analyses further elucidate the differences between cell types, where similarity is higher among cell types with homologous lineages, as illustrated in Supplementary Figure 8b.

**Figure 4.**
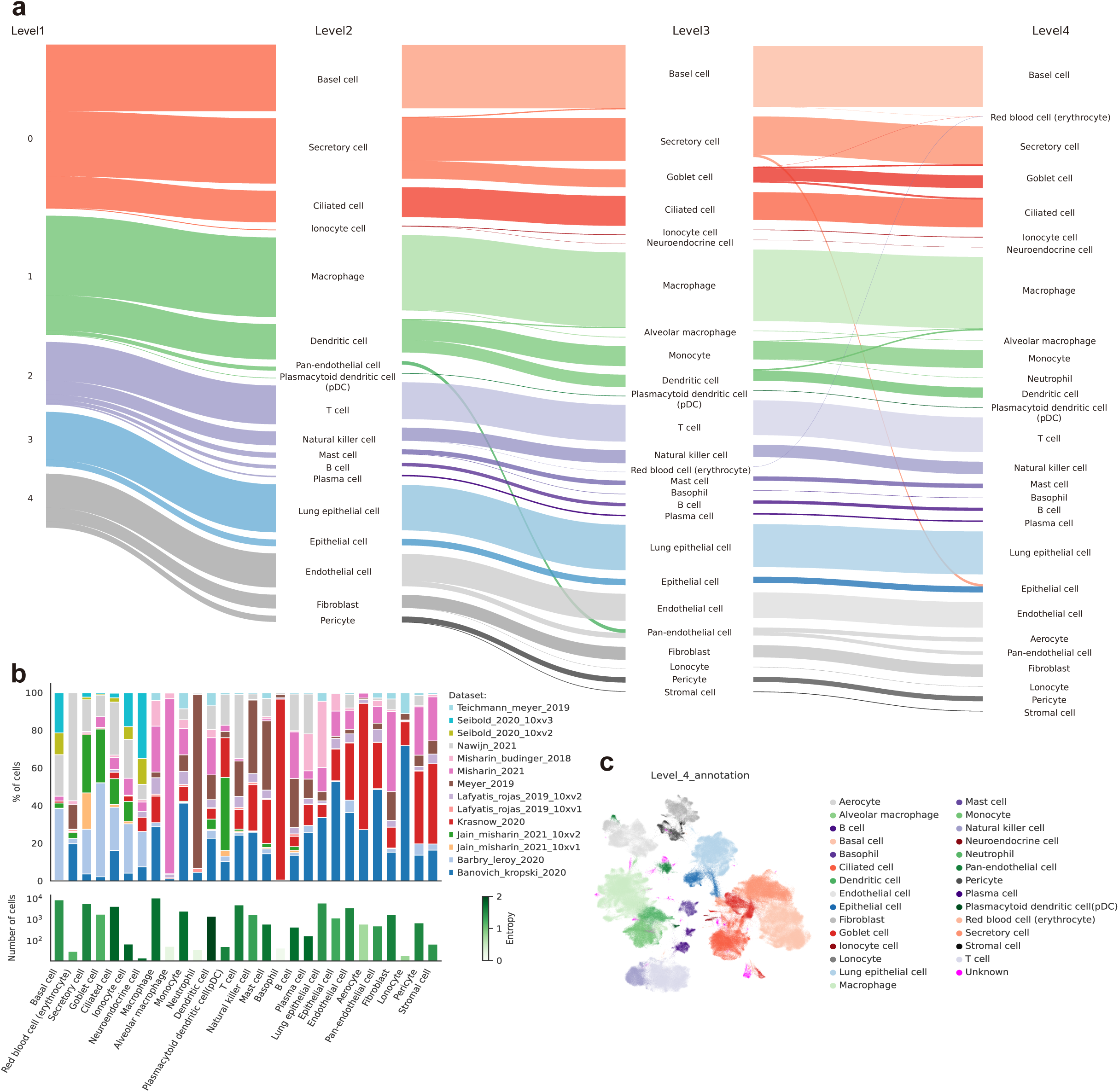
**a. Visualizations of hierarchical cell type annotations by Decoupler.** Bio-Copilot employs the Decoupler tool to annotate the cell clusters from level 2 to level 4. **b. The diversity and entropy of original independent datasets for each well-annotated cell type.** The majority of cell types comprise cells from multiple datasets. **c. UMAP visualization of the re-annotated cell types in level 4.** A total of 4,579 cells are classified as unknown cell types at level 4.

Furthermore, we further compare the differences between the cell types re-annotated by Bio-Copilot and the original cell types, as illustrated in Supplementary Figure 9a-c. It can be found that there is a consistent granularity between the re-annotated cell types and the original cell types in level 2 (Supplementary Figure 9a) and level 3 (Supplementary Figure 9b). Although the cell type nomenclature is different, the two cell type labels of most cells are from homologous lineages and can be well aligned. The re-annotated cell types in level 4 (Supplementary Figure 9c) have lower granularity resolution than the original cell types, which is limited by the cell type nomenclature of the database of human cell marker genes used by Bio-Copilot. Besides, we further analyzed unknown cell types at each level. As the level of clustering increases, more subclusters are classified as unknown cell types, as illustrated in Supplementary Figure 7b and Figure 4c. Finally, a total of 4,579 cells are classified as unknown cell types at level 4, as illustrated in Figure 4c. They are all isolated from major cell populations, and are used for exploratory research of rare cell types in downstream analysis task.

More details of Bio-Copilot performing the cell type annotation task refer to the Experimental Setup section in Methodology.

#### Downstream analysis

Following the cell type annotation task, Bio-Copilot performs downstream analysis task to explore new scientific discoveries, including rare cell type analysis and variance between individuals explained by covariates.

Bio-Copilot first analyzes the unknown cell types with a total of 4,579 cells. By analyzing the relationship between the unknown cell types from level 2 to level 4, there are three unknown cell types in the level 2 (i.e., ‘0_5’, ‘1_5’ and ‘3_3’), and their subtypes can be traced to these three unknown cell types, as illustrated in Supplementary Figure 10a-b. The other unknown cell types in level 3 and level 4 can be traced to well-annotated cell types. Therefore, we merge the unknown cell subtypes upward and keep the unknown cell types in the upper levels, as illustrated in Figure 5a. Subsequently, Bio-Copilot calculated the distribution of cells within each unknown cell type across the original independent datasets, alongside entropy values, as illustrated in Figure 5b. Obviously, most unknown cell types comprise cells from individual dataset and the number of cells is relatively small. Therefore, these unknown cell types can be classified as rare cell types. Bio-Copilot further analyzes the three rare cell types derived from level 2, i.e., ‘0_5’, ‘1_5’ and ‘3_3’. The original cell type distributions for these rare cell types are illustrated in Figure 5c and Supplementary Figure 10 c-d. Obviously, the rare cell types ‘1_5’ and ‘3_3’ were annotated as macrophage and alveolar type II in original dataset, respectively. The outlier characteristics of rare cell types ‘1_5’ and ‘3_3’ indicate that they may be cell subtypes of macrophage and alveolar type II, with specific gene expression and cell functions. The rare cell type ‘0_5’ has more original cell types, that can be traced back to epithelial cells, as illustrated in Figure 5c. However, this rare cell type has a significant difference from the epithelial cell, as illustrated in Figure 5a. Therefore, Bio-Copilot performs differential expression analysis at level 2 to obtain specifically expressed genes for these three rare cell types. Then, Bio-Copilot calls Enrichr^115,116^ enrichment analysis tool to parse the biological processes of each rare cell type, as illustrated in Figure 5d. The results of enrichment analysis suggest biological processes that are significantly activated or inhibited, and can guide the design of further biological experiments, such as validating the function of key genes, probing regulatory networks, or developing new therapeutic strategies.

**Figure 5.**
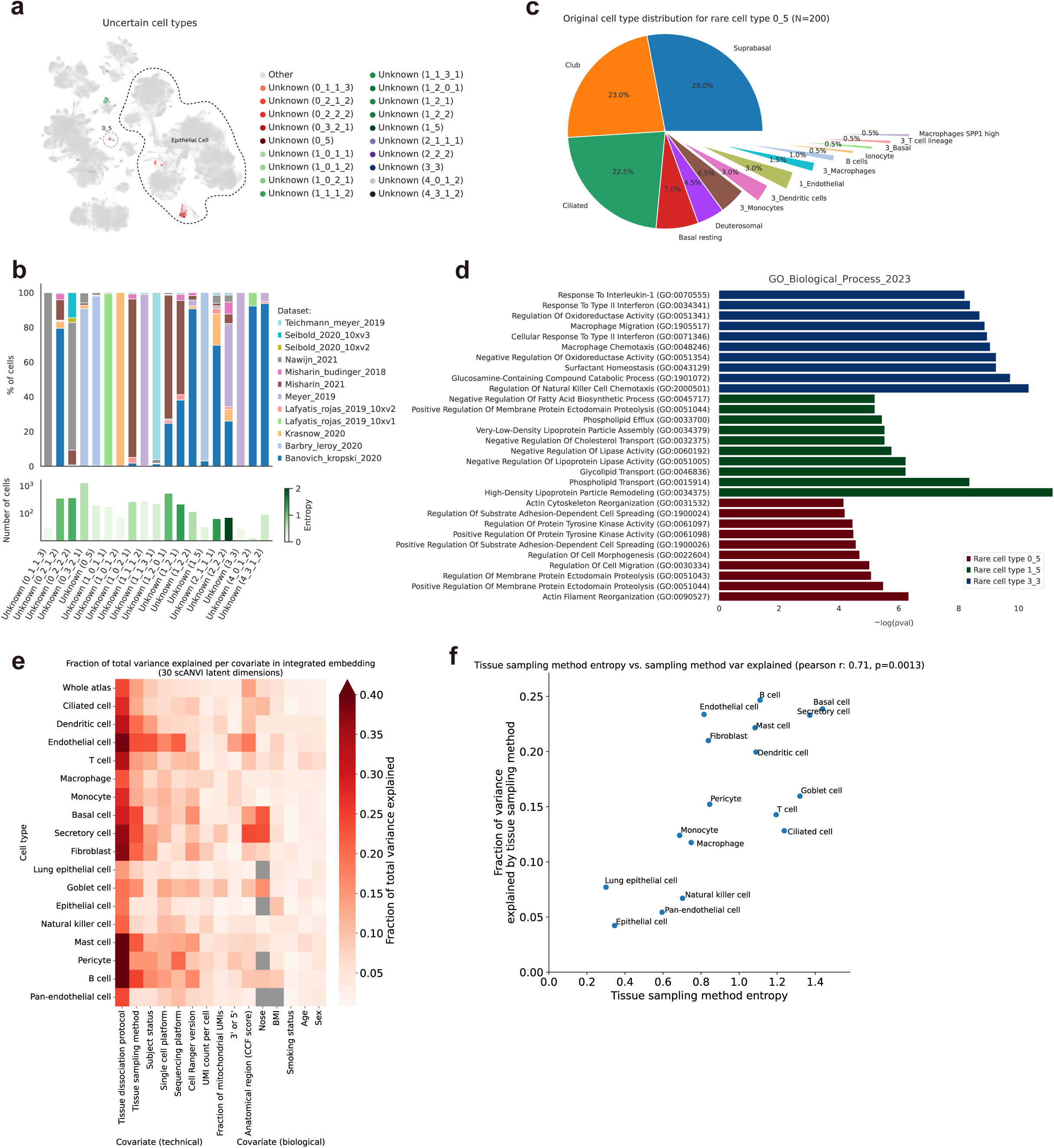
**a. UMAP visualization of unknown cell types annotated by Decoupler.** Most unknown cell types are located in outliers. **b. The diversity and entropy of original independent datasets for each unknown cell type.** Most unknown cell types comprise cells from individual dataset and the number of cells is relatively small. **c. The original cell type distribution of ‘0_5’.** The rare cell type ‘0_5’ can be traced back to epithelial cells. **d. The enrichment results of typical rare cell types.** Bio-Copilot employs Enrichr enrichment analysis tool to parse the biological processes of each rare cell type. **e. Variance explained per covariate across different cell types.** Bio-Copilot employs the embeddings of scANVI to conduct variance analysis. **f. Variance explained by tissue sampling methods versus their entropy.** Cell types that incorporate a greater variety of tissue sampling methods display more significant variation, with a positive correlation coefficient.

While the data integration task effectively mitigates batch effects, residual batch effects persist. Bio-Copilot employs the embeddings of scANVI to conduct variance analysis across different cell types and covariates, with results illustrated in Figure 5e. The variance analyses can reveal that technical covariates, such as tissue dissociation protocol, tissue sampling method, and single cell platform, still demonstrate noticeable variation contributions across many cell types. Specifically, tissue sampling methods such as ‘brush’, ‘biopsy’, ‘donor_lung’, ‘scraping’, and ‘surgical_resection’ show varied contributions to differential expression across cell types, as illustrated in Figure 5e. Notably, cell types that incorporate a greater variety of tissue sampling methods display more significant variation, with a positive correlation coefficient of 0.71, as illustrated in Figure 5f. Biological covariates, in contrast, exhibit lower contributions to variation across different cell types. Some cell types belonging to epithelial cells have relatively higher biological differences, such as secretory cell, goblet cell, basal cell and ciliated cell, as illustrated in Supplementary Figure 10e. It shows that they are less affected by technical covariates. For cell types sampled in fewer than 40 samples, the mean variance explained across all covariates showed a high negative correlation with the number of samples, as illustrated in Supplementary Figure 10f. This also underscores that adopting uniform technical protocols can significantly alleviate batch effects. Variance analyses conducted by Bio-Copilot yields conclusions consistent with those presented in the original study.

## IV. Discussion

### A. Summary

In the future trend of data-intensive and computation-intensive in big biological data processing and knowledge-intensive and intelligence-intensive in the convergence of interdisciplinary, this study proposes a data-intelligence-intensive Bio-Copilot system to facilitate hypothesis-free exploratory research and inspire novel scientific insights in large-scale omics. Bio-Copilot forms high-quality intensive intelligence through close collaboration between multiple agents, driven by LLMs, and human experts. We devise an agent group management strategy, an effective human-agent interaction mechanism, a shared interdisciplinary knowledge database, and continuous learning strategies to augment the capabilities of Bio-Copilot. In the benchmark test, Bio-Copilot achieves the overall state-of-the-art performance in all tasks against GPT-4o, AutoBA and AutoGen, while showcases high execution efficiency.

Furthermore, in the application of constructing a large-scale human lung cell atlas, Bio-Copilot not only reproduces the processes detailed in the original study but also employs a hierarchical annotation strategy to illustrate the continuous nature of cellular states and explore the characteristics of rare cell types. These experiments demonstrate the capabilities of Bio-Copilot to perform data-intelligence-intensive complex bioinformatics research. This study also reveals the potential of integrating AI capabilities with expert knowledge to accelerate impactful biological discoveries and explore uncharted territories in life sciences.

### B. Limitations

Although Bio-Copilot exhibits enormous promise in tackling data-intelligence-intensive research tasks, there remain limitations and areas requiring further development. Competition among agents is conducive to the enhancement of Bio-Copilot capabilities. Multiple agents can gain new knowledge through competition with other agents, serving as a strategy to continuous learning. For example, multiple agents perform the same task at the same time, use the evaluation function to evaluate the results of each agent, and then compare the results of each agent to obtain knowledge, and screen excellent solutions. Multiple agents usually require a large amount of computing resource overhead, especially when dealing with complex tasks and large-scale data, how to use these resources efficiently remains a critical challenge. Promising solutions include dynamic resource allocation, elastic computing, containerization, and microservice architectures that dynamically increase or decrease compute resources based on demand to improve resource utilization efficiency. Furthermore, complex tasks usually need to be decomposed into sub-tasks parallel or data parallel, and how to maintain load balance and control energy consumption and cost is also a key problem. Task prioritization scheduling algorithms and load balancing mechanisms are promising strategies to address these complexities. Additionally, incorporating energy-efficient management strategies, designing more efficient algorithms, leveraging heterogeneous computing resources, and employing model compression techniques can significantly reduce the cost of computation and energy consumption. Efficiently collecting, transmitting, processing, storing, and managing data to accurately monitor task progress and resource utilization poses another critical challenge. Current promising solutions entail lightweight data acquisition, data compression, streaming data processing, distributed data storage, and dashboard visualizations for real-time insights.

Designing effective prompts^117^ for user interaction with multiple agents, ensuring clear communication and efficient task delegation, presents a critical challenge. Vague, inconsistent, overly complex prompts, or those lacking coherent context, can lead to misdirected agent decisions and erroneous outputs, highlighting the importance of precise and contextually appropriate prompting strategies. Current prompt engineering approaches include: utilizing concise, clear, and neutral prompts; adopting stepwise guidance, incremental processing, and conversational prompts for complex tasks; reusing context, implementing positive feedback loops, and incorporating knowledge retrieval mechanisms to mitigate comprehension deviations. Furthermore, enhancing the diversity and adaptability of interactions to varying user needs and contexts poses a critical challenge. This includes supporting multifaceted interaction modes such as voice, vision, and text, alongside adaptive learning algorithms and feedback mechanisms to meet personalized interaction requirements. Potential solutions include multimodal fusion, cross-modal learning, online learning, user profiling.

### C. Potential future improvements

With the rapid advancement of AI technologies, AI agents have demonstrated substantial potential across various domains and ushered in new development opportunities. The integration of AI agent technology with cutting-edge technologies such as embodied intelligence^118,119,120^, world models^121^, and digital twins^122^ continues to drive innovation and advancements in various fields.

Embodied intelligence serves as a physical extension of AI agents into the real world, which can manipulate the senses and act of robots to interact with physical environment, and combine physical experience with cognitive processes. AI agents focus on autonomous learning, decision-making, and task execution based on data and algorithms, while embodied intelligence emphasizes physical interaction capabilities. When AI agents integrate embodied intelligence, they gain enhanced capacity to manipulate the physical environment, which requires the agent to have strong cognitive processing and physical interaction capabilities. For example, medical robots use the cognitive abilities of AI to diagnose patients and then perform specific physical therapies, embodying the synergy of intelligence and physical interaction. Embodied intelligence also has broad prospects in life science research, which can bring revolutionary changes to precision medicine, automated analysis of omics data, cell and molecular experiments, and ecological environment research.

The world model is the cognition and abstraction of AI agent’s environment, which can enhance the agent’s ability to perceive and understand changes in environmental states, and improve decision-making and action capabilities. Integrating AI agents with world models enables them to better understand and adapt to complex, dynamic environments. For example, in robotic navigation and autonomous driving, world models representing roads, traffic, and pedestrians enable agents to make decisions and navigate, avoiding obstacles and identifying optimal routes. In life sciences, world models apply to simulating disease spread, drug discovery, studies in biological evolution and genetics, integrating diverse data from organisms, compounds, and physical environments, thereby empowering agents with heightened environmental perception, informed decision-making, and precise actions.

Digital twins serve as virtual replicas of physical entities or environments, facilitating bidirectional communication between the physical realm and its digital counterpart. The convergence of AI agents with digital twins amplifies their capabilities and adaptability, encompassing real-time data analytics, simulation-based optimization, fusion of multi-source data, and environmental simulations. In smart cities, digital twins supplying real-time traffic data empower AI agents to optimize signal timing through refined decision-making, effectively alleviating congestion. In life sciences, digital twins integrate multi-source biological data and furnish virtual environments that enable AI agents to perform numerous simulations, thereby diminishing risks associated with physical experiments and expediting the research process.

In the future, AI agents will transcend their roles as mere human assistants to become collaborative partners, augmenting human capabilities across diverse tasks and delivering more intelligent decision-making support. AI agents will facilitate broader and more convenient applications in domains such as autonomous driving, smart factories, precision medicine, drug discovery, and scientific research, seamlessly integrating into daily life and work routines.

## V. Methodology

### A. Technical specifications of Bio-Copilot

#### Overall framework

The overall architecture of Bio-Copilot comprises multiple agents, a coordination layer, and an environment, as illustrated in Figure 1b. Agents, as the core components of Bio-Copilot, utilize LLMs as their central control nucleus. Based on task allocation, agents with different roles undertake distinct responsibilities. The coordination layer is responsible for mediating interactions between agents and the environment, as well as orchestrating the behaviors of agents. The environment, as an external context of Bio-Copilot, influences the behavior of agents, while the actions of these agents in turn modify the state of the environment.

Bio-Copilot employs AutoGen as its underlying framework that enables developers to customize task-specific agents from the predefined agent modules, and incorporates natural language conversation protocols to facilitate interactions between users and agents. In the coordination layer of Bio-Copilot, we design an agent group management strategy in the form of community structure to administer agents in various roles and coordinate inter-group communications. This strategy is optimized for addressing large-scale bioinformatics research tasks involving the engagement of numerous agents. Specifically, the task of constructing a large-scale human lung cell atlas employs three agent groups: coarse-level planning, fine-level planning, and action execution. Additionally, we devise three effective strategies within the coordination layer: A user-friendly and efficient human-agent interaction mechanism supports close collaboration between human experts and multiple agents; A shared interdisciplinary knowledge database enhances the cognitive capabilities of agents through knowledge empowerment, and supports task-driven extraction of relevant information; Continuous learning strategies, such as self-reflective learning, enhance the decision-making and execution capabilities of agents, thereby rapidly accommodating emerging and challenging bioinformatics research tasks.

#### Agent group management strategy

We illustrate the agent group management strategy using the case of agent groups engaged in the task of constructing a large-scale human lung cell atlas. Each group comprises multiple agents fulfilling distinct roles, and those agents sharing the same role across different groups exhibiting slightly different variations in responsibilities.

##### Coarse-level planning group

As the name suggests, the coarse-level planning group aims to formulate a coarse-grained task execution plan based on user inputs that typically comprise task specifications and data descriptions. This group has three agents: ‘Planner’, ‘User Proxy’ and ‘Reviewer’. ‘Planner’ initially receives user inputs and generate an overall plan. Subsequently, ‘Reviewer’ meticulously reviews the overall plan and propose s suggestions for revisions. Afterwards, ‘Planner’ revises the overall plan based on suggestions, and relays the revised overall plan to ‘User Proxy’. ‘User Proxy’ provides a static human-agent interaction interface, enabling users to verify whether the overall plan requires any further adjustments.

##### Fine-level planning group

The fine-level planning group has similar agent roles and internal communication protocols to the coarse-level planning group, consisting of ‘Refined Planner’, ‘User Proxy’, and ‘Reviewer’. The difference is that the fine-level planning group receives a single step from the overall plan devised by the coarse-level planning group as input, to generate and refine a more detail plan for that step.

##### Action execution group

Action execution refers to code generation, execution and debugging, involving data loading and writing, information retrieval, and tool invocation. This group consists of four agents including ‘Coder’, ‘Executor’, ‘User Proxy’ and ‘Checker’. Specifically, ‘Coder’ receives the detail plan for the current step from the fine-level planning group and generates the corresponding code. ‘Executor’, equipped with a Python kernel and a code execution environment, is responsible for receiving and executing the code generated by ‘Coder’. Following code execution, ‘Checker’ receives outcomes returned by ‘Executor’, undertakes code debugging, identifies issues within the current code, suggests remedial modifications, and returns these to ‘Coder’ as feedback. ‘User Proxy’ is responsible for the termination. Moreover, within the action execution group, ‘User Proxy’ offers two types of human-agent interaction interfaces (i.e., static and dynamic). The static interface is exposed following the initial receipt of code by ‘Coder’, so that users can inspect the code for any potential issues before execution. The timing of the dynamic interface exposure is determined by ‘Checker’. ‘Checker’ will instruct ‘User Proxy’ to expose the dynamic human-agent interface in three specific situations. Firstly, ‘Checker’ receives a repeated execution error, indicating ‘Coder’ inability to correct the error. Secondly, ‘Checker’ receives more than five error messages pertaining to code anomalies. Thirdly, ‘Checker’ identifies instances necessitating manual specification of names within errors, such as actual gene names present in the dataset. Additionally, the code generation capability of ‘Coder’ is enhanced through continuous learning by feeding its output from the previous step into the current one, ensuring the continuity and consistency of generated code. When the code is executed successfully, ‘User Proxy’ will terminate the group conversation.

##### Communications between agent groups

With in each agent group, we have introduced how agents interact. In this section, we will comprehensively elaborate the communication mechanisms between agent groups. The three agent groups communicate approximately linearly. As illustrated in Figure 2, the coarse-level planning group decomposes user-assigned task into a multi-step overall plan following predefined rules, with subsequent refinements. Then, the fine-level planning group iteratively conducts further detailed planning on each step of the overall plan in a loop manner, subsequently refining the detailed execution plans for each step. The action execution group is responsible for executing the detailed execution plan of each step, involving code generation, running, and debugging. Upon completion of a step, the process iterates to the next one until the entire overall plan is accomplished.

#### Human-agent interaction mechanism

Although Bio-Copilot can autonomously carry out many routine bioinformatics analysis tasks, human-agent collaboration remains the most effective approach in tackling complex and dynamic tasks. There are three primary reasons: Firstly, bioinformatics analysis tasks often entail personalized data processing stages, where agents relying on their inherent knowledge may struggle to accommodate personalized requirements. Secondly, when agents cannot autonomously resolve encountering errors, they risk being locked in infinite loops, hindering progression to subsequent steps. Lastly, insufficient user input can lead to misinterpretations by agents due to information gaps, resulting in erroneous decisions or failed executions.

In the data-intelligence-intensive research paradigm, both human experts and AI agents serve as core participants in the scientific investigation. To support the tightly coupled human-agent collaborative workflow, we design a user-friendly and efficient human-agent interaction mechanism for Bio-Copilot. Departing from conventional static human-agent interaction models, where the timing of human interaction with Bio-Copilot is fixed, we devise a dynamic human-agent interaction mechanism that enables agents to determine the timing of exposing the human-agent interaction interfaces based on the fulfillment of predetermined conditions, thereby ensuring high real-time interaction capabilities. This dynamic human-agent interaction mechanism is extensively employed within the action execution group, that can enhance the capabilities of agents to tackle complex and dynamic tasks. Bio-Copilot integrates both dynamic (appears in action execution group) and static (appears in all agent groups) human-agent interaction mechanisms, optimizing autonomous task execution while minimizing unnecessary human intervention.

#### Shared interdisciplinary knowledge database

We establish a shared interdisciplinary knowledge database within Bio-Copilot, thereby augmenting the cognitive capacities of agents with diverse roles through knowledge empowerment. By collecting and encoding knowledge from multidisciplinary research articles into a form comprehensible to agents, the shared interdisciplinary knowledge database enables agents to retrieve relevant information. For instance, during coarse-level planning, ‘Planner’ initiates its process by retrieving relevant information from the knowledge database, which is then strategically integrated as supplementary cues into its planning workflow. Knowledge-driven insights augment the input of ‘Planner’, thereby the planning quality of ‘Planner’ is enhanced by enriching the decision-making process with informed perspectives. In Bio-Copilot, knowledge empowerment is crucial for achieving high autonomy and efficiency in handling complex biological data and tasks, which can alleviate biases and illusions inherent in reasoning and decision-making processes.

#### Continuous learning

In order to rapidly accommodate emerging and challenging bioinformatics research tasks, we devise continuous learning strategies for Bio-Copilot, incorporating self-reflective learning. In Bio-Copilot, agents engage in self-reflective learning by reviewing their own behaviors, progressively refining autonomous decision-making and execution capabilities over time, thereby enhancing task efficiency in subsequent missions through process streamlining, selecting optimal tools, and minimizing manual intervention. For example, the self-reflective learning mechanism is embodied in two kinds of agents, i.e, ‘Reviewer’ and ‘Checker’, that are responsible for examining the plans formulated by ‘Planner’ (or ‘Refined Planner’) and the code produced by ‘Coder’. Through iterative interactions and self-reflective learning, they have refined their capabilities to autonomously suggest modifications, thereby improving the quality of feedback.

### B. Experimental Setup

#### Datasets

Data for constructing the core HLCA (human lung cell atlas) is derived from 11 independent studies^123,124,125,126,127,128,129,130,131,132^, integrating 14 datasets comprising 168 health lung tissue samples from a total of 107 individual donors, with diversity in age, sex, ethnicity, BMI and smoking status. The HLCA project has preprocessed these datasets, involving harmonization of cell type nomenclature, alignment of gene nomenclature, data quality control, and data normalization. Specifically, the HLCA project established a five-level hierarchical cell type nomenclature to harmonize cell types across datasets, aligned gene nomenclature to the GRCh38^133^ human genome reference with Ensembl^134^ standardization, applied quality control to exclude cells with fewer than 200 genes and genes expressed in less than 10 cells, and performed SCRAN^135^ normalization on raw counts for uniformizing UMI counts per cell.

#### The workflow of constructing a large-scale lung cell atlas

The workflow of constructing a large-scale human lung cell atlas is accomplished through a collaborative effort between researchers and Bio-Copilot, which mainly comprises four tasks: dataset integration task, hierarchical clustering task, cell type annotation task, and downstream analysis task. For each task, researchers specify the task objectives, prompting Bio-Copilot to generate an initial execution plan, which then iteratively refined via extensive human-machine interactions, guided by researcher feedback informed by insights and experience. Additionally, during the task execution phase of Bio-Copilot, researchers can actively engage in human-machine interactions to identify and rectify incorrect actions of the system.

##### Data integration

To integrate disparate single-cell datasets of lung tissue, data integration algorithms are employed to yield an integrated embedding that aligns cells of the same type across independent datasets within a feature expression space. This unified embedding facilitates the construction of a comprehensive and unbiased cell atlas, enabling insightful analyses into the biological diversity and heterogeneity within lung tissue. Bio-Copilot queries single-cell data integration methods to get current existing numerous approaches, such as BBKNN^136^, fastMNN^137^, Harmony^138^, Scanorama^139^, scGEN^140^, scANVI^111^, scVI^141^, and Seurat^142^. Through further document retrieval, Bio-Copilot gained knowledge from the research study of scIB^143^ benchmarking, that evidences the capability of scANVI in achieving the optimal comprehensive performance. Hence, Bio-Copilot employs scANVI to perform single-cell dataset integration, and retrieves its code example for automatic code generation. Next, we describe the key parameter settings in the order of code, all of which are set by Bio-Copilot and can be adjusted according to the feedback of experts. To prepare the input of the top 2,000 highly variable genes (HVGs) for scANVI, the scanpy.pp.highly_variable_genes function provided by the scanpy^144^ toolkit is employed to process raw count data, configured with parameters: batch_key=’dataset’, min_mean=0.6, flavor=’cell_ranger’, n_top_genes=2000, leaving default settings for all other parameters. Furthermore, scANVI can use cell labels to improve performance, thus the curated original cell types are employed. Guided by user prompts, Bio-Copilot utilizes the names of the independent study datasets as batch-covariates, sets the number of hidden features to 30 for scANVI, while other parameters are automatically configured to their defaults: n_layers=2, encode_covariates=True, deeply_inject_covariates=False, gene_likelihood=’nb’, with GPU acceleration enabled via use_cuda=True. Employing GPUs significantly accelerates the training process of the scANVI method. The trained scANVI model outputs the hidden embeddings of the integrated datasets, which have undergone batch-effect correction and are utilized for subsequent clustering analyses and visualization tasks.

##### Hierarchical clustering

Following data integration, the integrated dataset comprises 587,218 cells. To gain deeper insights into cellular diversity and heterogeneity, Bio-Copilot performs a hierarchical clustering with 4 levels to delineate cell populations by iterating from coarse to finer resolutions. Firstly, the K-nearest neighbor graph of the hidden embeddings is constructed using the scanpy.pp.neighbors function, with the parameter for the number of neighbors set to n_neighbors=30. At the first level of clustering (i.e., the coarsest level), a leiden clustering function scanpy.tl.leiden is employed to generate the first level cell population clusters, with the clustering resolution parameter of resolution=0.01. Subsequently, new K-nearest neighbor graphs for each cluster from the first level cell populations are constructed using the hidden embeddings and the scanpy.pp.neighbors function with n_neighbors=30 again. Reapplying leiden clustering to each cluster from the first level, with a clustering resolution parameter of resolution=0.1, produces the second level cell populations. Next, sequentially proceeding the third and fourth level analysis like second level ultimately generates a hierarchical, four-level structure of cell populations. In order to track relationships among cell populations effectively, hierarchical clusters are labeled according to their parental lineage and ordinal position among sisters, such that Cluster 1_2 denotes the third sub-cluster (counting from 0) of Cluster 1. Besides, Bio-Copilot employs scanpy.tl.rank_genes_groups function with default parameters in scanpy toolkit to perform differential expression analysis for clusters at each level. Furthermore, for visualizing cell clusters at each level, the K-nearest neighbor graph of the hidden embeddings is fed to the scanpy.tl.umap function for dimensionality reduction followed by scanpy.pl.umap function to plot 2D UMAPs, where the color parameter in scanpy.pl.umap is set to the level key of hierarchical clustering and the palette parameter defines the colors for cell clusters.

##### Cell type annotation

Bio-Copilot performs cell type annotation for cell clusters in the hierarchical structure. Considering the coarse granularity of first level clusters renders their annotation less informative, thus the cell type annotation is performed from level 2 to level 4. Bio-Copilot ingests a database of human cell marker genes from CellMarker^145^, and subsequently invokes the Decoupler tool to identify cell types for each cluster based on that database. Decoupler uses the over representation analysis (ORA) function to process a table for screening lung tissue associated cell types from the database and the integrated dataset with 2,000 HVGs, leaving default settings for all other parameters. Then, Decoupler employs statistical test function dc.rank_sources_groups to identify which are the top predicted cell types for each cluster, with parameters of groupby=’cluster_level_2’, reference=’rest’, method=’t-test_overestim_var’ for level 2 cell clusters. Based on the results of the statistical test, Bio-Copilot considers the rank of significance score and mean foldchange value to calculate a comprehensive ranking, annotating the top 1 cell type for each cluster. If a cluster has a top 1 significance score of less than 30, the cluster is classified as unknown cell type. This same strategy is then replicated for the annotation of level 3 and level 4 cell clusters, with parameters groupby=’cluster_level_3’ and groupby=’cluster_level_4’ in statistical tests, respectively. After completing the cell type annotation, Bio-Copilot perform some basic statistical analysis for each well-annotated cell type at level 4, including the number of cells, original dataset distribution, and information entropy. Furthermore, Bio-Copilot performs differential expression analysis for these well-annotated cell types at level 4 to uncover marker genes. The scanpy.tl.rank_genes_groups function in scanpy toolkit is utilized for differential expression analysis, with the groupby parameter set to the group of level 4 cell types, n_genes=2 specifying the top 2 differentially expressed genes to retrieve, and the parameter of statistical test method=’wilcoxon’. To refine the selection for more significant differentially expressed genes, the scanpy.tl.filter_rank_genes_groups function is applied for additional filtering with default parameters: min_in_group_fraction=0.7, max_out_group_fraction=0.2, min_fold_change=2. After filtering, the differentially expressed genes are ranked based on significance and inter-group variation, enabling the selection of a subset of top 2 genes as marker genes for each cell type.

##### Downstream analysis

Following the cell type annotation, Bio-Copilot performs rare cell type analysis and variance between individuals explained by covariates in downstream analysis to explore new scientific discoveries. For the rare cell type analysis, Bio-Copilot merge the unknown cell subtypes upward to keep the unknown cell types in the higher granularity, enabling top-down analysis. Bio-Copilot first performs basic statistical analysis for unknown cell types, including the number of cells, original dataset distribution, and information entropy. Based on these results, their rare properties are inferred. Bio-Copilot further analyzes the three significant rare cell types derived from level 2, including the original cell type distributions, and differential expression analysis with all cell types at level 2. The scanpy.tl.rank_genes_groups function in scanpy toolkit is utilized for differential expression analysis, with the groupby parameter set to the group of level 2 cell types, and the parameter of statistical test method=’wilcoxon’. The specifically expressed gene set for each rare cell type is obtained with significance P-value<0.05. Based on the specifically expressed gene set, Bio-Copilot calls Enrichr enrichment analysis tool to parse the biological processes using the GO_Biological_Process_2023 database. For variance between individuals explained by covariates, Bio-Copilot performs statistical analysis to elucidate the correlations between covariates in technical methods and biological characteristics. A quantitative assessment of the variance in gene expression profiles attributable to each covariate within every cell type is performed. To ensure robust statistical analysis, the cell types represented by fewer than two samples or those with a total cell count below 50 are excluded, while any sample within a cell type containing less than ten cells is also excluded. Initially, the covariate statuses and 30 scANVI latent embeddings per sample for every cell type are statistically summarized to generate matrices. Statistical summaries for scANVI latent embeddings, UMI counts, and mitochondrial ratios involve averaging across all cells within a sample, whereas other covariates are sampled from any cell within the sample, as they remain invariant among cells from the same sample. Within matrices derived for each cell type, linear regressions between each covariate and each scANVI latent embedding are conducted to predict the embedding feature explained by each covariate, subsequently calculating variances. Finally, the variances across all scANVI latent embeddings are summed to quantify the differential impact of covariates across individual samples. Bio-Copilot further calculates the correlations between variance explained and various covariates, including the correlation between the diversity in tissue sampling methods for a cell type and the fraction of variance explained by sampling method for that cell type, the correlation between the ratio of variance explained by biological covariates versus technical covariates for a cell type and the number of samples in that cell type, and the correlation between the mean variance explained by covariates in a cell type and the number of samples in that cell type.

#### Implementation Details

##### Computing resource

All analyses presented in the study run on Bio-OS^146^ with elastic resources, at least 128Gb RAM memory, 128 cores of 2.5GHz CPU, and a GPU with 32Gb memory. And the following python (v3.11) packages support for Bio-Copilot and lung cell atlas workflow are required: python==3.11.7, chainlit==1.0.100, autogen==1.0.16, pyautogen==0.2.19, openai==1.14.0, jupyter==1.0.0, httpx==0.23.0, jupyterlab==4.0.12, notebook==7.0.8, torch=1.7.1+cu110, scanpy==1.10.0, scarches^147^==0.3.5, decoupler==1.6.0, numpy==1.26.4, pandas=2.2.2, scikit-learn==1.4.2

##### Details of the system

The preliminary input from the user includes the task and dataset descriptions, separated by a line break. Upon receipt of this preliminary input, Bio-Copilot is initiated into operation. Specifically, the user’s task requirements are subjected to the coarse-level planning group to obtain a high-level plan that is used to extract steps with a step extractor. Then, each step is sent to the fine-level planning group for a detailed plan. Moreover, at the end stage of the coarse-level planning and the fine-level planning groups, Bio-Copilot will automatically provide an interactive interface for users to determine whether the plan completed by the agents still needs to be modified. The action execution group receives information from the fine-level planning group and is responsible for generating, debugging, and executing code. During the code debugging process, if a code error arises that the group cannot resolve independently, the group will provide an interactive interface for users to participate in solving the error. Once the code for the current step runs successfully, the action execution is terminated. Bio-Copilot cyclically executes fine-level planning and action execution groups according to the steps division until all are completed. When Bio-Copilot completes the user’s task, it exits the operational mode, and returns to a waiting state.

#### Details in the construction of the lung cell atlas

Data integration. Firstly, user provides the task and dataset descriptions as the initial input, that is, "Perform single-cell data integration. xxx.h5ad: all single-cell data need to be integrated and analysis for constructing the human lung cell atlas. The adata column of "integration_label" contains the label information for integration.". The main steps of coarse-level planning include: (i) Load and preprocess the data; (ii) Integrate the data using scANVI; (iii) Generate integrated embeddings; (iv) Save the integrated data. Next, when performing the data preprocessing step, the agent is prompted to omit the visualization step, "do not need to do any visualizations", to improve the running efficiency. When executing the scANVI model training step, the ‘Coder’ agent will call existing documents to assist in code writing, which uses the default parameter values in documents. The user can let the agent modify the corresponding parameter values according to the actual data situation and use the GPU to run the program, that is, "use os to set gpu id as 4, set unlabeled_category as ‘unlabeled’". Finally, when saving the integrated data, the agent is prompted to save the scANVI integration results of highly variable genes to the original adata, i.e., "save the integration result of scanvi to the original adata instead of adata_hvg".

Hierarchical clustering. User provides the task and dataset descriptions, that is, " Perform multi-level clustering of the cells in the HLCA. Note that counting the initial clustering, the total number of clustering levels is 4. For the final HLCA, you also need calculate marker genes for every clustering. xxx.h5ad: the integrated single-cell data with scANVI that needed to be multi-level clustered.

The integrated representation is in "X_scANVI" of the adata.". The main steps of coarse-level planning include: (i) Load the dataset; (ii) Initial clustering (level 1); (iii) Subsequent clustering (levels 2-4); (iv) Calculate marker genes for each cluster; (v) Save the final annotated dataset. When hierarchical clustering is performed, sub-cluster labels need to be generated and saved to adata and csv files for subsequent analysis. Therefore, this prompt is provided, "generate new sub-clustering labels for each cluster and update dataset with new sub-cluster labels.". When calculating marker genes for each cluster, the agent needs to be prompted to provide the groups parameter in sc.tl.rank_genes_groups according to the cluster label. In addition, two perspectives "sister" and "all" are adopted to calculate marker genes, and the difference between them needs to be suggested to the agent, i.e., "all is a standard computational logic. sister refers to the situation where when the cluster depth is greater than 1, you need to use the clusters within groups as adata to calculate marker genes.". Besides, parameters such as resolution used in clustering can also be artificially specified.

Cell type annotation. Similarly, user provides the task and dataset descriptions, that is, " Perform cell types annotation with python packages. xxx.h5ad: stores 3 level clustering label (cluster_level_2, cluster_level_3, cluster_level_4) which is used for annotation. highly_variable parameter can be used for selecting highly variable genes.". The mainly steps of the coarse-level planning include: (i) Load the necessary libraries for analysis; (ii) Load and preprocess the data; (iii) Prepare the database of marker genes for Decoupler; (iv) Run Decoupler for cell type annotation; (v) Save the annotated data. During the planning phase, human intervention removes unnecessary steps, "Delete the step that Initialize the Decoupler model with the default cell marker information. please annotation on all cluster levels.". In addition, when multiple agents cannot identify data structures or some personalized data processing is required, users also need to intervene through human interaction interfaces in a timely manner. For example, in the step of preparing the database of marker genes for Decoupler, provides "Set adata.var_names to ‘gene_symbols’ in adata.var" or "Retain data where adata.var.highly_variable is true for cell annotation and copy it.", and similar instructions.

Rare cell type analysis in downstream analysis. Firstly, user needs to describe the task and dataset, i.e., "I now have clustering and annotation results. I want to discover rare cell types. What should I do? I will provide you with all the data and information as follows. hlca_decoupler_annotation.h5ad: Already annotated cell data. The multi-level clustering results are stored in ‘cluster_level_1’, ‘cluster_level_2’, ‘cluster_level_3’, ‘cluster_level_4’.". The main steps of coarse-level planning include: (i) Load the annotated cell data; (ii) Identify and validate rare cell types; (iii) Validate rare clusters using annotation; (iii) Further analysis to confirm rare cell types; (v) Save the results. In the planning stage, user needs to guide the agent to conduct further analysis on possible rare cell types, providing guidance such as "You need to do further analysis to confirm rare cells, rather than simply judging by the existing clustering and annotation results." or "The current clustering and annotation results are all multi-level results. Can you use this information to do further analysis?" When confirming possible rare cell type clusters, the agent sets a threshold by default, and the user need to guide the specific threshold setting.

Variance explained by covariates in downstream analysis. User provides the task and dataset description as the initial input, i.e., “Determine how much of the variance in the integrated HLCA embedding is explained by each of the metadata covariates including You need to use the results of the fourth level annotation in ‘ann_level_4_dc_final’ for analysis. Note that your analysis should be conducted from sample-level observations.” The main steps of coarse-level planning include: (i) Load the data, (ii) Extract and aggregate relevant data, (iii) Perform variance analysis, (iv) Save the model results. Since the task contains a lot of personalized data processing and conditional judgments, the user needs to provide relevant prompts. For instance, in the data load and preprocessing steps, unnecessary processing is eliminated, and the prompt “do not need to handle missing values” is given. When integrating data from the sample level (step (ii)), Bio-Copilot will save all dimensions of "X_scANVI" into one parameter, which makes subsequent variance analysis impossible. Therefore, the user must provide the prompt as "Store each component of the X_scanvi_embed as a separate row in the aggregated_row. You can first fill other data into aggregated_row. Then use a for loop to add component keys into aggregated_row with changing key names". In variance analysis (step (iii)), users need to give a prompt like "Y may be NAN or string value, you should deal with these conditions.". Besides, users can provide necessary parameter settings to Bio-Copilot. Otherwise, Bio-Copilot will automatically set parameter values or generate code according to default values.

### C. Evaluation Metrics

#### Metrics for Bio-Copilot benchmarking

##### Task planning evaluation

This section aims to verify whether Bio-Copilot can achieve superior planning performance compared to control group and ablation group. We have selected the following 10 metrics to measure the performance of different methods:

- Clarity. It evaluates the explicitness and logical coherence of the generated plan in its expression. A clear plan should allow the executor to understand and implement it without ambiguity.
- Technical Feasibility. It considers the practical operability of the solutions proposed in the plan. Technical feasibility includes assessing the necessary resources, current technological limitations, and potential risks.
- Structurality. The structurality metric focuses on the organization and layout of the plan. Good structurality implies that the plan has a clear beginning, middle, and end phases, with logical coherence between each stage.
- Comprehensiveness. The comprehensiveness metric measures whether the plan has considered all relevant constraints and objectives. A comprehensive plan should cover all aspects of the issue without omitting key execution steps or objectives.
- Rationality of Method Selection. It assesses the match between the chosen algorithm and the needs of the plan. A rational choice of algorithm should be based on the problem’s characteristics and constraints to ensure the plan’s efficiency and effectiveness.
- Risk Management. Assesses the identification, analysis, and mitigation strategies for potential risks and uncertainties in the plan.
- Data Management. Refers to systematic organization, storage, and handling of data to ensure its accuracy, accessibility, and security throughout the research process.
- Scientific Rigor. Involves the strict application of scientific methods and principles to ensure the reliability, validity, and reproducibility of research findings.
- Resource Allocation. Examines the efficiency and appropriateness of the resources (time, budget, personnel, equipment) allocated for the task.
- Scalability. Evaluates the potential for the plan to be scaled up or adapted for larger or different contexts if successful.

We use GPT-4 to score the above indicators for task planning evaluation in all tasks, where the prompt is illustrated in Supplementary Figure 1a.

#### Code quality evaluation

This section aims to validate the effectiveness of Bio-Copilot in generating reliable and executable task code. We have adopted the following 12 metrics to measure and compare the performance of different methods:

- Accuracy. Assesses whether the generated code can accurately implement the established functions and requirements.
- Efficiency. It focuses on the performance of the code in terms of resource consumption, including but not limited to execution time, memory usage, and CPU utilization rate.
- Completeness. Measures whether the generated code includes all necessary components to achieve the intended functionality.
- Readability. Evaluates the clarity and understandability of the code, involving the appropriateness of naming conventions, structural organization, and comments.
- Logicality. It checks the coherence and rationality of the internal logical flow of the code, including the correct application of conditional statements, loop structures, and exception handling.
- Portability. Evaluates the code’s ability to be transferred between different environments or datasets.
- Maintainability. This dimension assesses how easily the code can be modified or extended in the future. It includes factors like modularity, use of comments, and adherence to coding standards.
- Reusability. This dimension evaluates the extent to which parts of the code can be reused in different projects or contexts. It looks at the generality and modularity of the code.
- Robustness. This dimension assesses the code’s ability to handle unexpected inputs or conditions without crashing or producing incorrect results. It includes error handling and validation mechanisms.
- Security. This dimension assesses the code’s ability to protect against unauthorized access, data breaches, and other security threats. It includes secure coding practices and data protection measures.
- Usability. This dimension evaluates how user-friendly the code is, particularly for non-developers or domain experts. It includes aspects like clear documentation, intuitive interfaces, and ease of use.
- Scalability. This dimension measures how well the code can handle increasing amounts of data or more complex computations without significant performance degradation.

We also use GPT-4 to score the above indicators for code quality evaluation in all tasks, where the prompt is illustrated in Supplementary Figure 1b.

#### Execution efficiency evaluation

This section aims to evaluate the task completeness, user interventions, runtime, computing resource cost of Bio-Copilot during task execution.

- Task completeness. The proportion of completed steps to the total number of steps.
- User interventions. The total number of conversations initiated by users, each of which is counted as one human-agent interaction round.
- Runtime. The total time from the start of a task until completion or termination.
- Computing resource cost. Average CPU, GPU, and memory usage during task execution.

#### Comprehensive score

We use the following formula to synthesize task planning evaluation, code quality evaluation, and execution efficiency evaluation to calculate a comprehensive performance score for each method,

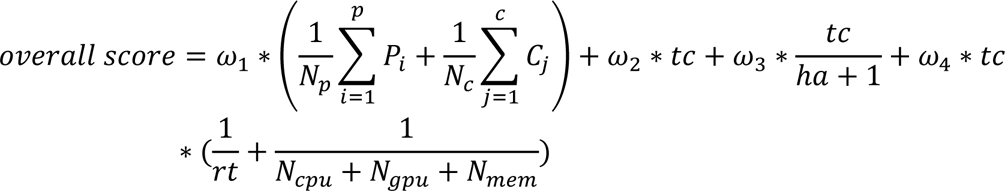

where *ω_1_*∼*ω_4_* indicate the weights, *N_p_* = 10 indicates the 10-evaluation metrics for task planning, *N_C_*= 12 indicates the 12-evaluation metrics for code quality, *tc* indicates task completeness, *ℎa* indicates the number of user interventions, *rt* indicates the runtime of a task, *N_Cpu_*, *N_gpu_*, *N_mem_* indicate average CPU, GPU, and memory usage during task execution. In this study, the weights are set as *ω_1_* = 2, *ω_2_* = 20, *ω_3_* = 5, *ω_4_* = 1 for the comprehensive performance score, as illustrated in Figure 3a.

### Metrics for lung cell atlas analysis

#### Entropy

We employ Shannon entropy to quantify the amount of uncertainty or information content inherent in a discrete distribution. Formally, it measures the average amount of information conveyed by each event in a discrete probability distribution. The Shannon entropy is defined as

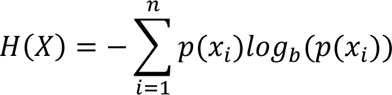

where *n* is the number of possible outcomes, *p*(*xi*) represents the probability of the *itℎ* outcome occurring, and the base of the logarithm *b* determines the unit of measurement (commonly, *b* = 2 for bits). A higher entropy value suggests higher uncertainty or disorder; conversely, a lower entropy implies more predictability or order within the system. In this study, Shannon entropy helps in identifying cell type label disagreement for a specific level of annotation, and calculating the diversity of donors or tissue sampling methods in a cluster.

## VI. Data availability and Code availability

Data and code are available in https://doi.org/10.5281/zenodo.11210015 and https://github.com/lyyang01/Bio-Copilot respectively.

## Acknowledgements

This work was supported by self-supporting Program of Guangzhou National Laboratory (Grant No. SRPG22007), youth science foundation of Guangzhou National Laboratory (Grant No. QNPG23-12).

## Author contributions

Conceptualization: Yixue Li. Supervision: Jiao Yuan, Yixue Li.

Algorithm development and implementation: Yang Liu, Lu Zhou, Rongbo Shen.

Datasets collection, processing and application: Rongbo Shen, Yang Liu, Lu Zhou.

Methods comparisons: Yang Liu, Lu Zhou, Rongbo Shen.

Biological interpretation: Rongbo Shen.

Manuscript writing and figure generation: Rongbo Shen, Yang Liu, Lu Zhou.

Manuscript reviewing: Yixue Li.

All authors approved the manuscript.

---

**Supplementary Figure 1.**
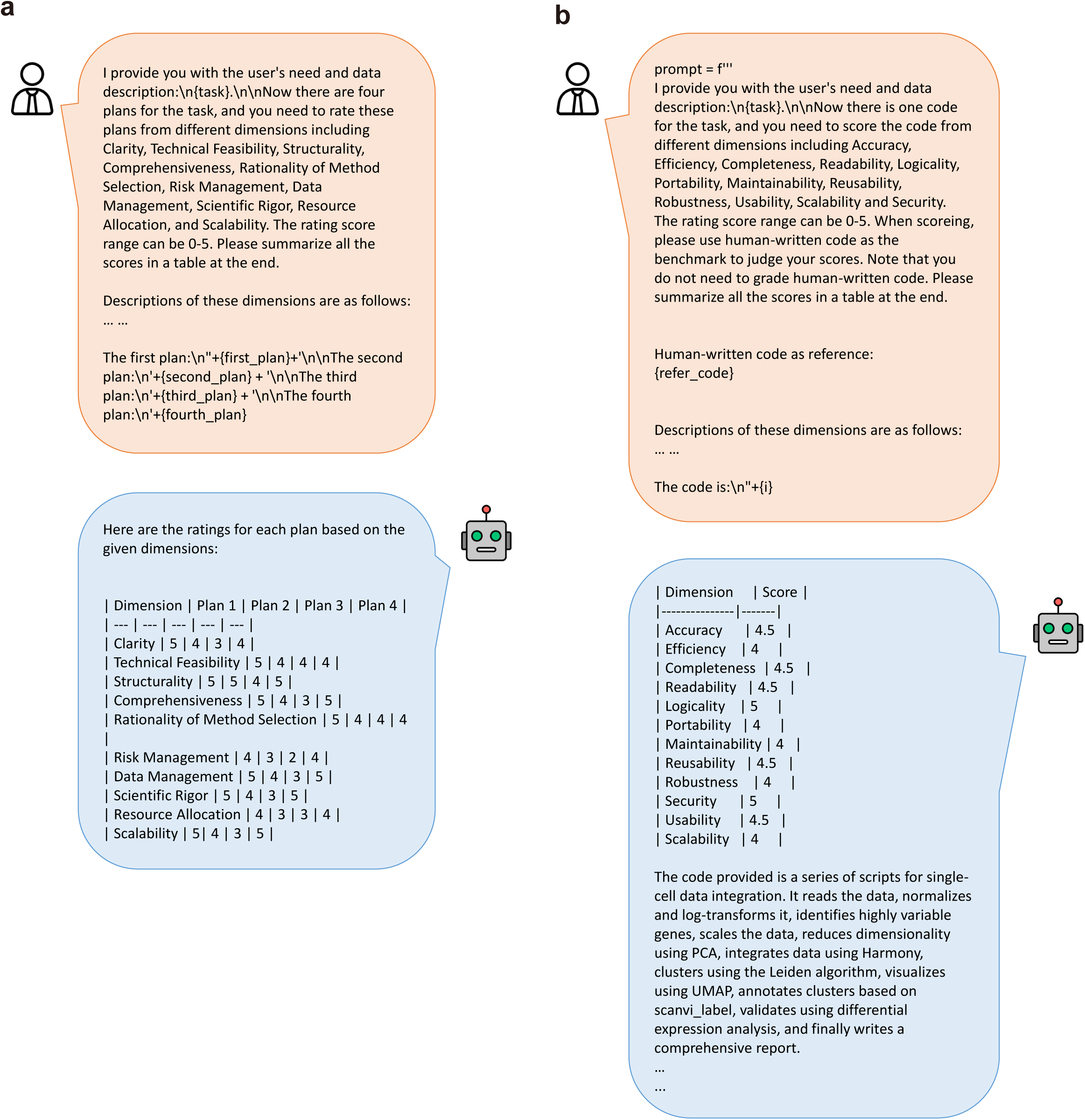
**a. The prompt of GPT-4 to score task planning. b. The prompt of GPT-4 to score code quality.**

**Supplementary Figure 2.**
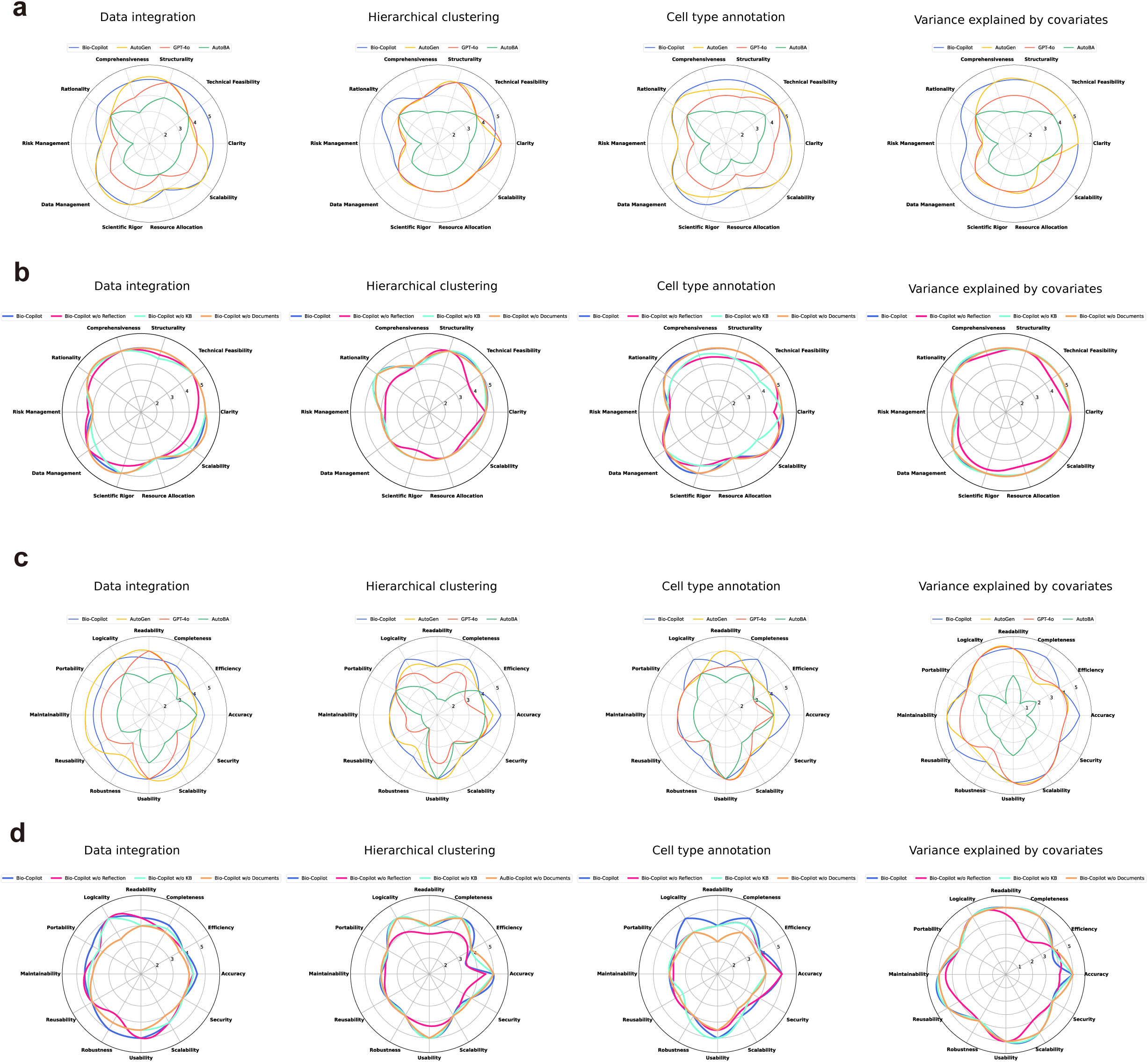
**a. The scores of task planning in control group. b. The scores of task planning in ablation group. c. The scores of code quality in control group. d. The scores of code quality in ablation group.**

**Supplementary Figure 3.**
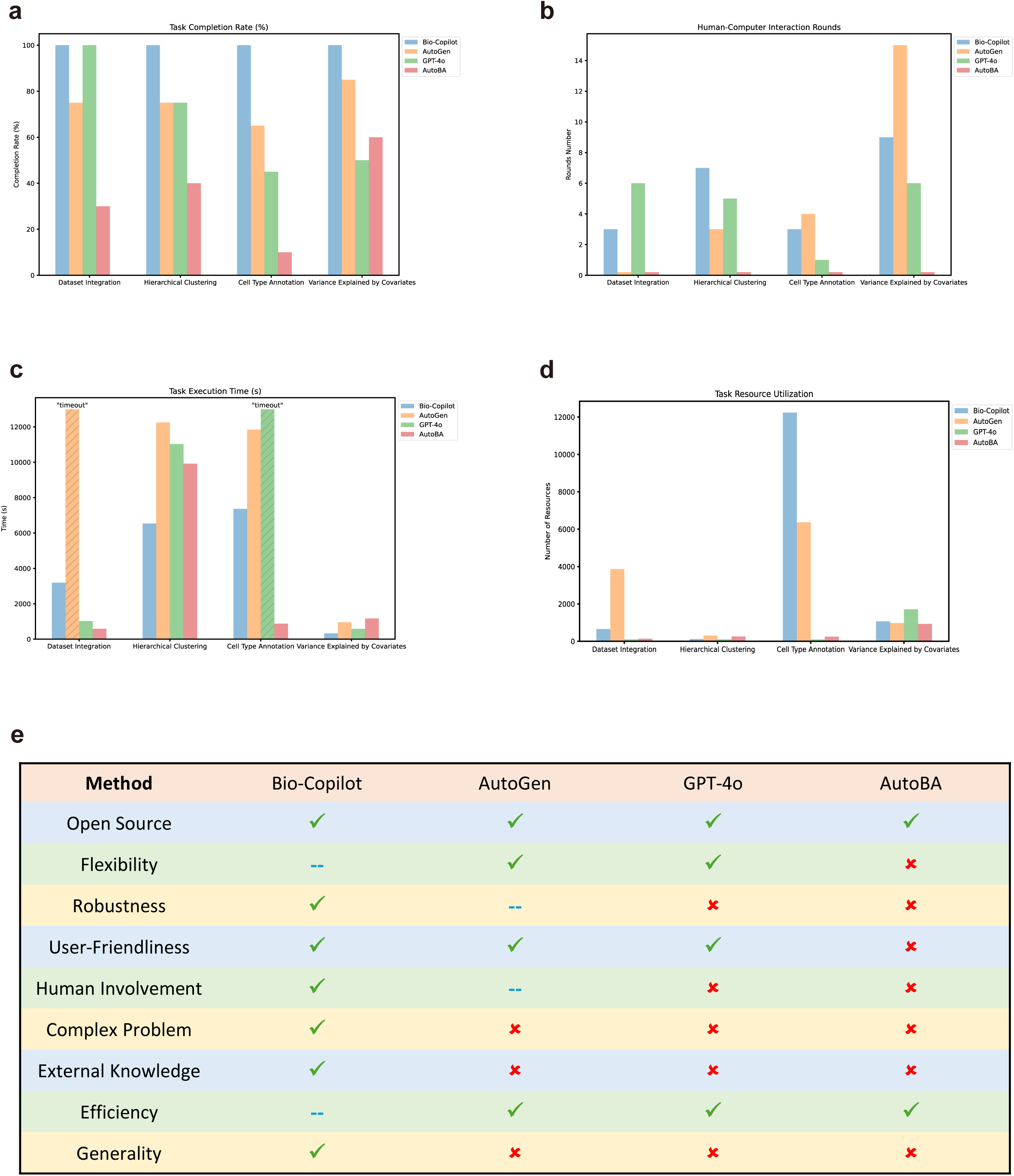
**a. The performance of task completeness. b. The performance of user interventions. c. The performance of runtime. d. The performance of computing resource cost. e. Features of methods.**

**Supplementary Figure 4.**
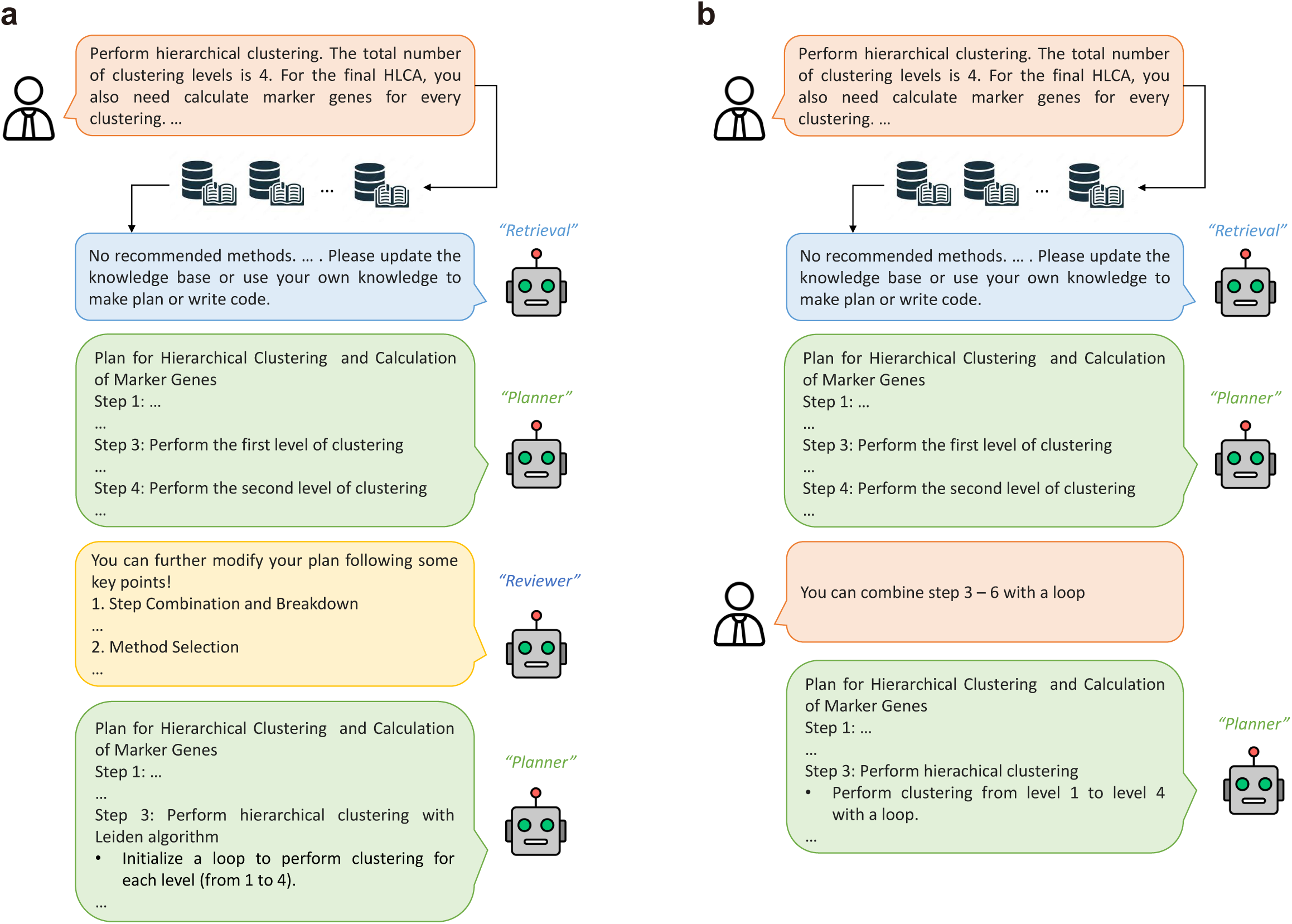
**a. The workflow of Bio-Copilot with self-reflective learning. b. The workflow of Bio-Copilot without self-reflective learning.**

**Supplementary Figure 5.**
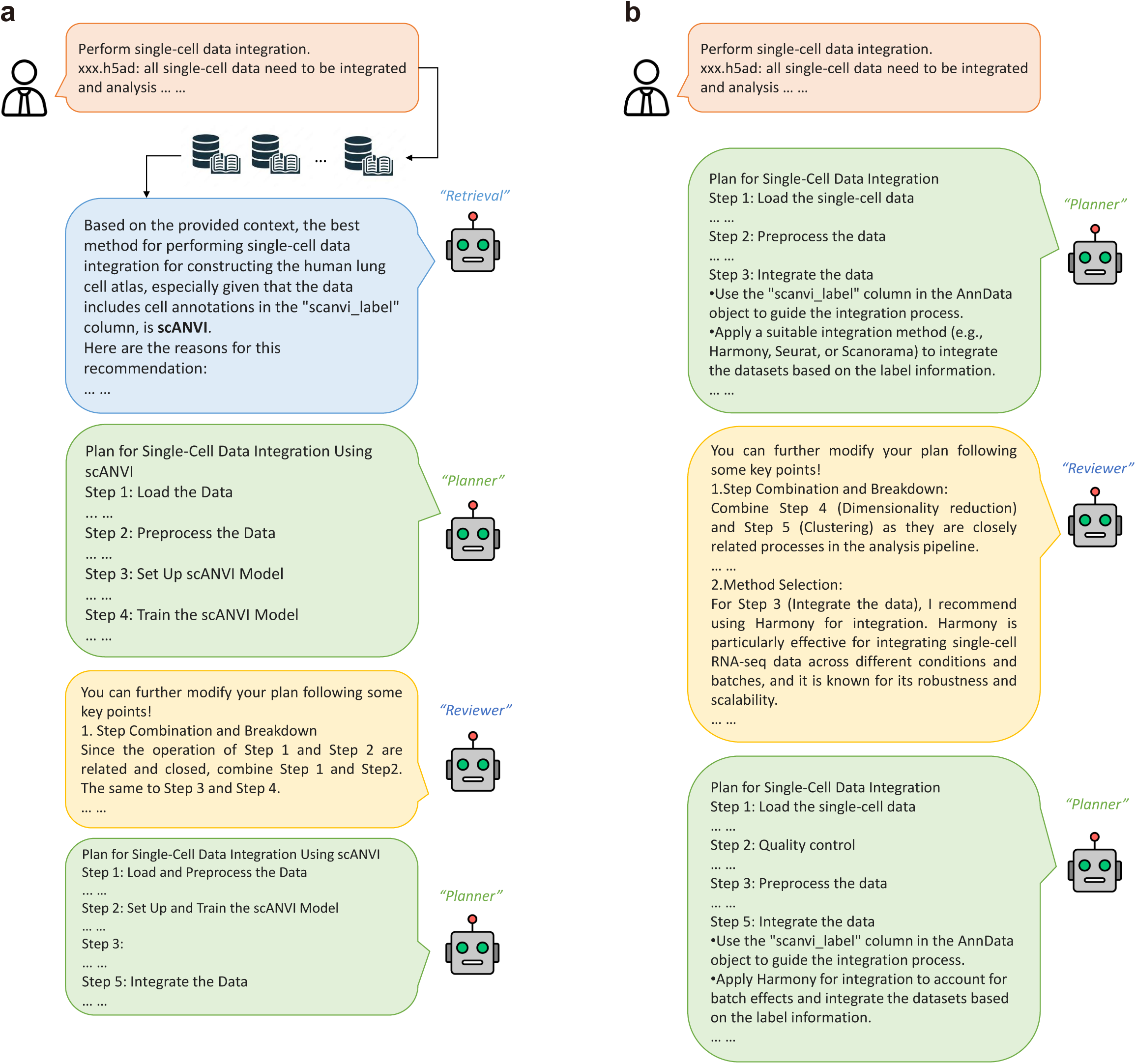
**a. The workflow of Bio-Copilot with knowledge empowerment. b. The workflow of Bio-Copilot without knowledge empowerment.**

**Supplementary Figure 6.**
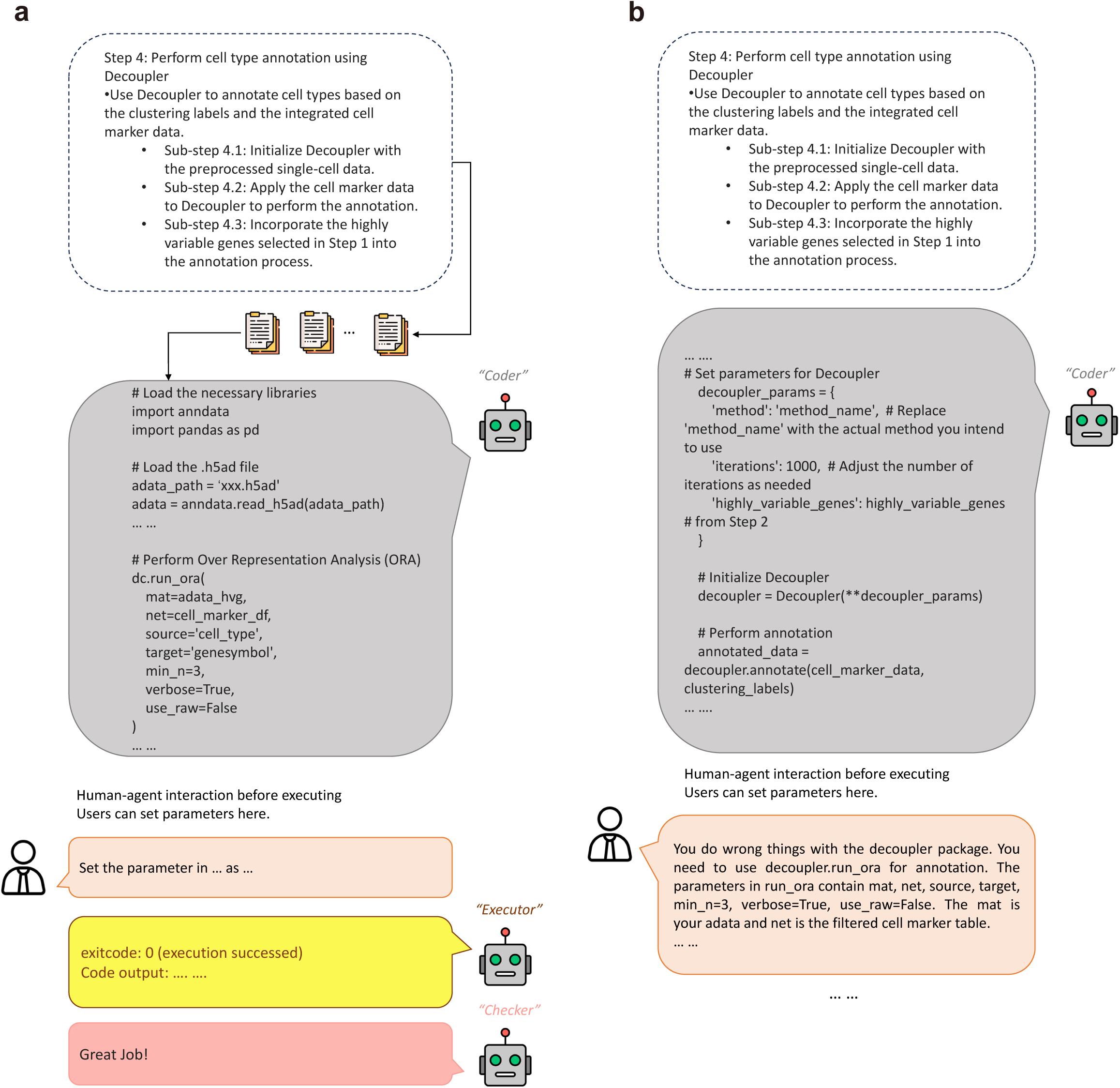
**a. The workflow of Bio-Copilot with document retrieval. b. The workflow of Bio-Copilot without document retrieval.**

**Supplementary Figure 7.**
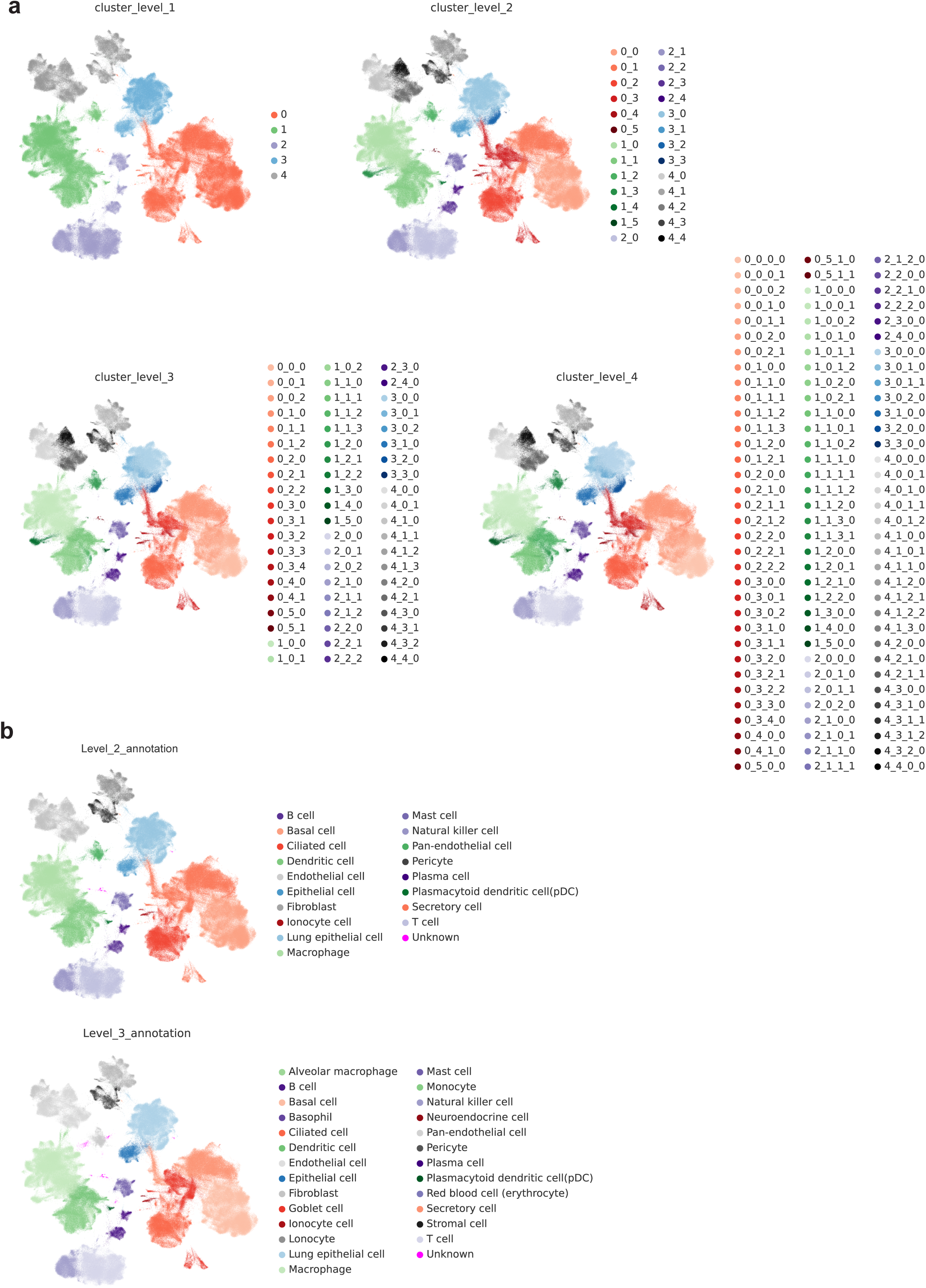
**a. UMAP visualizations of the hierarchical clustering results. b. UMAP visualizations of the re-annotated cell types in level 2 and level 3.**

**Supplementary Figure 8.**
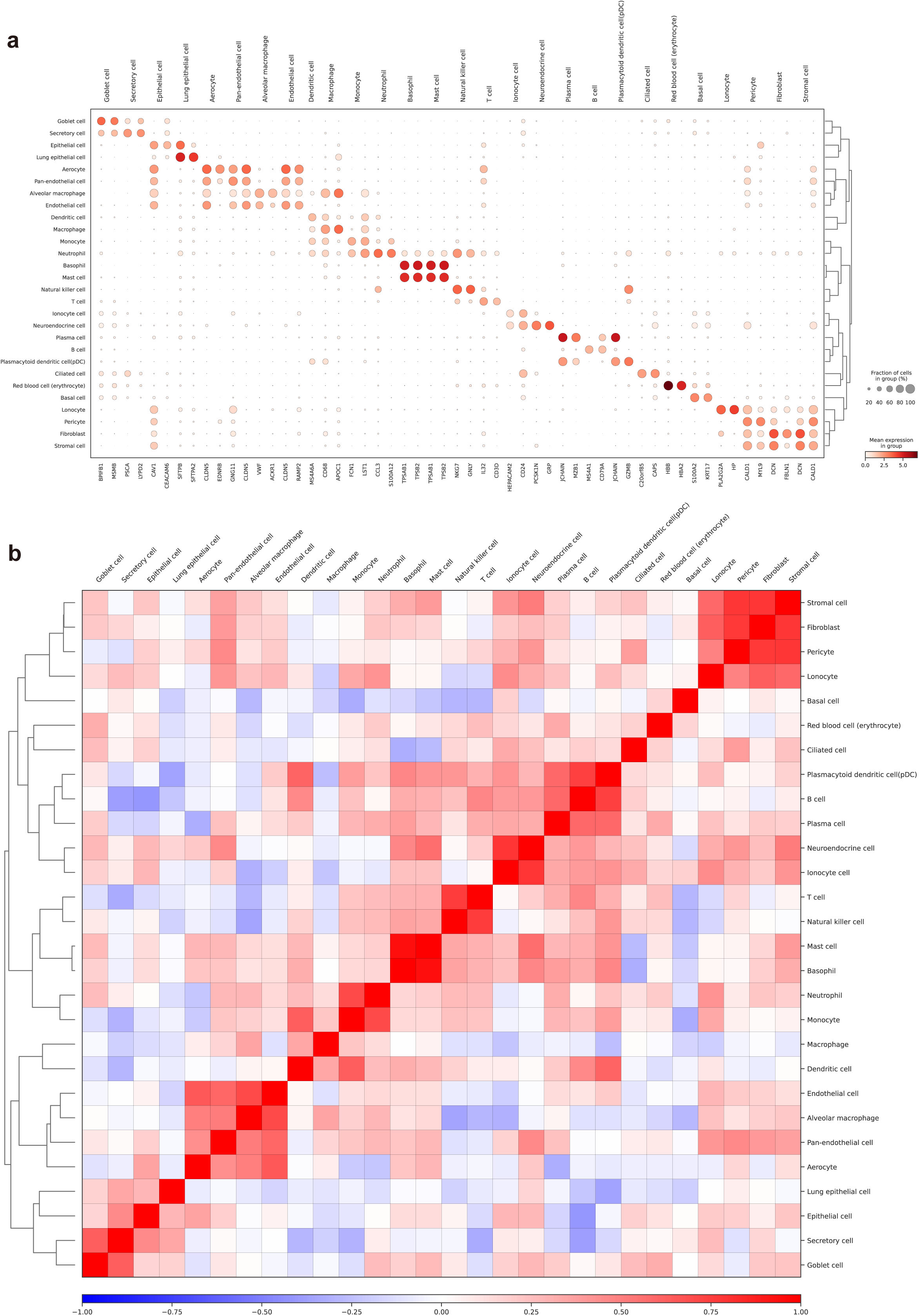
**a. Differential expression analysis of well-annotated cell types. b. Correlation between well-annotated cell types.**

**Supplementary Figure 9.**
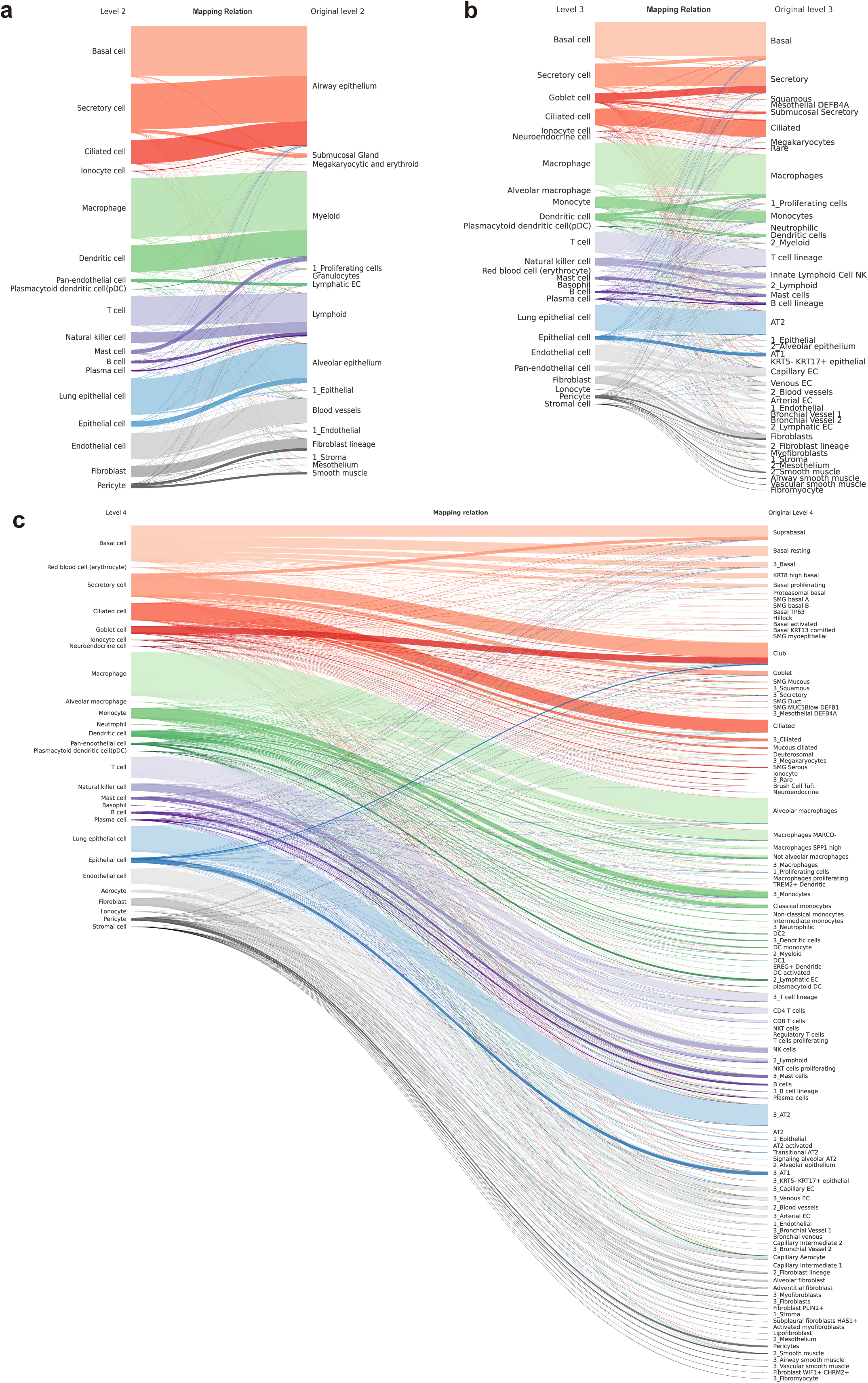
**a-c. Differences between the cell types re-annotated by Bio-Copilot and the original cell types at level 2, 3, and 4, respectively.**

**Supplementary Figure 10.**
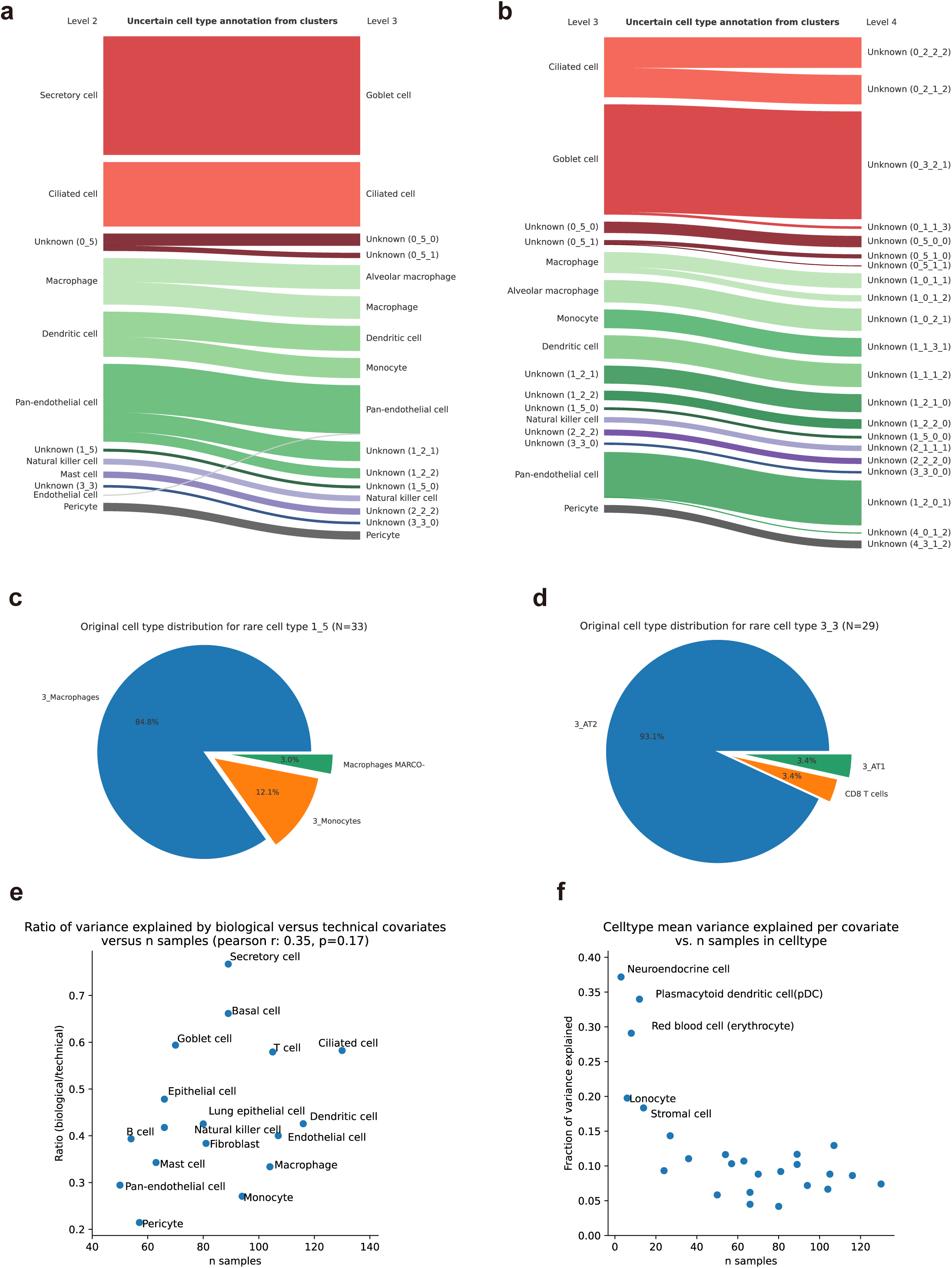
**a-b. Relationship between the unknown cell types from level 2 to level 4. c. The original cell type distribution of ‘1_5’. d. The original cell type distribution of ‘3_3’. e. Variance explained by biological versus technical covariates. f. Mean variance explained per covariate.**

